# Development of a multiplexed targeted mass spectrometry assay for LRRK2-phosphorylated Rabs and Ser910/Ser935 biomarker sites

**DOI:** 10.1101/2020.11.23.394486

**Authors:** Raja S. Nirujogi, Francesca Tonelli, Matthew Taylor, Pawel Lis, Alexander Zimprich, Esther Sammler, Dario R Alessi

**Author notes:** Correspondence to RSN or DRA.

## Abstract

Mutations that increase the protein kinase activity of LRRK2 are one of the most common causes of familial Parkinson’s disease. LRRK2 phosphorylates a subset of Rab GTPases within their Switch-II motif, impacting interaction with effectors. We describe and validate a new, multiplexed targeted mass spectrometry assay to quantify endogenous levels of LRRK2-phosphorylated Rab substrates (Rab1, Rab3, Rab8, Rab10, Rab35 and Rab43) as well as total levels of Rabs, LRRK2 and LRRK2-phosphorylated at the Ser910 and Ser935 biomarker sites. Exploiting this assay, we quantify for the first time the relative levels of each of the pRab proteins in different cells (mouse embryonic fibroblasts, human neutrophils) and mouse tissues (brain, kidney, lung and spleen). We define how these components are impacted by Parkinson’s pathogenic mutations (LRRK2[R1441C] and VPS35[D620N]) and LRRK2 inhibitors. We find that the VPS35[D620N], but not LRRK2[R1441C] mutation, enhances Rab1 phosphorylation in a manner blocked by administration of an LRRK2 inhibitor, providing the first evidence that endogenous Rab1 is a physiological substrate for LRRK2. We exploit this assay to demonstrate that in Parkinson’s patients with VPS35[D620N] mutations, phosphorylation of multiple Rab proteins (Rab1, Rab3, Rab8, Rab10 and Rab43) is elevated. We highlight the benefits of this assay over immunoblotting approaches currently deployed to assess LRRK2 Rab signalling pathway.

## Introduction

Autosomal dominant mutations that hyperactivate leucine rich repeat kinase 2 (LRRK2) are a frequent cause of late onset familial Parkinson’s disease [1, 2]. LRRK2 is a large multidomain enzyme made up of armadillo, ankyrin, leucine-rich repeats, tandem Roco type GTPase, Ser/Thr kinase domains, and a C-terminal WD40 domain [3, 4]. Recent tomography and cryo-EM analysis provide the first glimpse of the LRRK2 structure revealing how Parkinson’s mutations and inhibitors modulate the closed and open conformation of the kinase and impact upon microtubule binding [5, 6]. The best characterized substrates of LRRK2 comprise a subset of Rab GTPases, namely Rab3A, Rab3B, Rab3C, Rab3D, Rab8A, Rab8B, Rab10, Rab12, Rab29, Rab35 and Rab43 [7, 8]. LRRK2 phosphorylates these Rab proteins at a conserved Ser/Thr residue (Thr 72 for Rab8A and Thr73 for Rab10) located within the effector-binding Switch-II motif. In addition, LRRK2 efficiently phosphorylates Rab1 in vitro and in overexpression studies at the equivalent residue (Thr75) [8, 9], but thus far, we have not observed LRRK2 mediated phosphorylation of endogenous Rab1 in either wild type or pathogenic LRRK2 knock-in cells [7].

LRRK2-phosphorylated Rab proteins lose their ability to bind their cognate effector proteins, and instead interact with new sets of phospho-binding effectors such as RILPL1/2 and JIP3/4 [7, 10]. LRRK2 suppresses ciliogenesis in striatal cholinergic interneurons through LRRK2-phosphorylated Rab10 forming a complex with RILPL1 [11, 12]. Recent evidence also points towards LRRK2 controlling lysosomal and endomembrane homeostasis through its ability to phosphorylate Rab8A [13, 14]. Parkinson’s causing D620N autosomal dominant mutation in the VPS35, the cargo binding subunit of the retromer complex also elevates LRRK2 mediated Rab protein phosphorylation through an unknown mechanism [15].

Various, dominantly inherited pathogenic mutations within the Roc (N1437H, R1441G/C/H), Cor (Y1699C), and kinase (G2019S, I2020T) domains of LRRK2 have been well-characterized [16]. The G2019S mutation is located in the conserved Mg^2+^ subdomain VII motif of the kinase domain and represents the most commonly observed pathogenic mutation [17]. All LRRK2 pathogenic mutations enhance phosphorylation of Rab10 and several of the other Rab protein substrates studied, typically between 1.5 to 4-fold [7, 8, 18, 19]. Pathogenic mutations also stimulate LRRK2 autophosphorylation at Ser1292 [20]. However, stoichiometry of Ser1292 phosphorylation is extremely low, making it hard to detect and quantify robustly, especially for endogenous LRRK2 using available phospho-specific antibodies [21]. Standard mass spectrometry approaches are also rendered difficult as the tryptic peptide encompassing Ser1292 lies within a two amino acid phospho-peptide. LRRK2 is also phosphorylated on several well studied serine residues including Ser910 and Ser935, located between the ankyrin and leucine rich repeats that regulate 14-3-3 binding [22]. These sites become rapidly dephosphorylated upon pharmacological inhibition of LRRK2 [23], and have thus been widely used to assess the target engagement of LRRK2 inhibitors [24]. Phosphorylation of Ser910 and Ser935 does not correlate with intrinsic LRRK2 kinase activity and moreover, several pathogenic mutations including R1441C/G suppress the phosphorylation of these residues through an unknown mechanism [22, 23].

Measurement of Rab protein phosphorylation is the gold-standard approach to readout the steady-state activity of endogenous LRRK2 pathway in cell or tissue extracts. Global mass spectrometry (MS) analysis points towards Rab8A and Rab10 comprising the most abundant LRRK2 Rab substrates in cells and tissues analysed [8]. Recent work employing a sensitive, targeted MS-based assay established that stoichiometry of Rab10 Thr73 phosphorylation is low, typically 1-3% [25]. It is likely that stoichiometry of other LRRK2-phosphorylated Rab proteins will be significantly lower. Most widely utilized current approaches to assess LRRK2 mediated Rab protein phosphorylation rely on antibody-based approaches. Thus far, selective phospho-Rab monoclonal antibodies have been developed that specifically detect pRab10 phosphorylated at Thr73 [26] and Rab12 phosphorylated at Ser105 [15]. In addition, a pan-selective phospho-antibody detecting Rab proteins phosphorylated on Thr residues (Rab1 Rab3, Rab8, Rab10, Rab35 and Rab43) has been described [26]. This pan-selective pRab antibody has been exploited to immunoprecipitate multiple LRRK2-phosphorylated Rab proteins that can then be detected by immunoblot or MS analysis [7, 27]. This antibody, however, does not detect LRRK2 substrates Rab12 or Rab29 that are phosphorylated on Ser residues [26].

Here, we describe a new multiplexed targeted Parallel Reaction Monitoring (PRM) [28]. MS assay designed to quantity multiple LRRK2-phosphorylated Rab proteins in a single assay. This assay permits simultaneous quantitation of endogenous LRRK2-phosphorylated Rab1, Rab3, Rab8, Rab10, Rab35 and Rab43 in 0.1 to 1 mg of cell or tissue extract. Our assay also quantifies total levels of Rab proteins and LRRK2 in addition to LRRK2-phosphorylated Ser910 and Ser935. Employing this assay, we are able to quantify the relative abundance of the different pRab proteins in multiple cells and tissues, defining the impact of LRRK2 inhibitors and Parkinson’s disease causing mutations.

## Materials and Methods

A comprehensive list of reagents, antibodies, cDNA constructs, mice, cell lines, buffers, equipment, software packages utilized in this study are provided in Supplementary file 1.

### Cell Culture, Treatments, and Lysis

MEFs were cultured in Dulbecco’s modified eagle medium (DMEM) supplemented with 10% (v/v) fetal calf serum, 2 mM L-glutamine, 100 U/ml penicillin, 100 μg/ml streptomycin, nonessential amino acids and 1 mM sodium pyruvate. All cells were grown at 37°C and 5% (v/v) CO2 in a humidified atmosphere. Cell lines utilized for this study were tested regularly for mycoplasma contamination and confirmed as negative prior to experimental analysis. Cells were lysed in ice-cold lysis buffer containing 50 mM Tris/HCl pH 7.4, 1 mM EGTA, 1 mM sodium orthovanadate, 10 mM 2-glycerophosphate, 50 mM NaF, 5 mM sodium pyrophosphate, 270 mM sucrose, supplemented with 1 μg/ml microcystin-LR, 1 mM sodium orthovanadate, complete EDTA-free protease inhibitor cocktail (Roche), and 1% (v/v) Triton X-100. Lysates were clarified by centrifugation for 20 min at 20,800 g at 4°C. The supernatant was collected in a new low binding Eppendorf-tubes (#022431081) and protein concentrations of cell lysates were determined using the Bradford assay. The lysates were flash frozen in liquid nitrogen and stored at −80 °C.

### Generation of mouse embryonic fibroblasts (MEFs)

Wildtype and homozygous MEFs were isolated from littermate-matched mouse embryos at day E12.5, as described in a previous study resulting from crosses between heterozygous mice [29]. Wild type and homozygous LRRK2 [R1441C] [7], VPS35[D620N] [15] and LRRK2 [G2019S] MEFs have been described previously.

### Neutrophil isolation

Neutrophils were isolated from peripheral human blood from healthy donors and Parkinson’s patients with VPS35[D620N] mutations as described previously [30]. Formal ethical approval for these studies was granted. Viability and cell lysis performed as described previously [30]. Briefly, Neutrophils were isolated from freshly drawn whole blood from 3 donors (25 ml per donor) using the EasySep Direct Human Neutrophil Isolation Kit and Easy 50 EasySep Magnet (Stemcell) as described [30, 31]. After isolation, neutrophils from each donor were resuspended in RPMI and treated with either the indicated concentration of MLi-2 or DMSO (0.1% v/v). After 1h, cells were harvested in 100 μl per sample of ice-cold lysis buffer supplemented with the potent protease inhibitor diisopropylfluorophosphate (DIFP) to a final concentration of 0.5 mM, that was added to lysis buffer just before lysing cells [31].

### Mice

Mice were maintained under specific pathogen-free conditions at the University of Dundee (U.K.). All animal studies were ethically reviewed and carried out in accordance with Animals (Scientific Procedures) Act 1986 and regulations set by the University of Dundee and the U.K. Home Office. Animal studies and breeding were approved by the University of Dundee ethical committee and performed under a U.K. Home Office project license. Mice were multiply housed at an ambient temperature (20–24°C) and humidity (45–55%) and maintained on a 12 h light/12 h dark cycle, with free access to food (SDS RM No. 3 autoclavable) and water. For the experiments described in Figure 9 (VPS35[D620N] knock-in model), in Figure 8 (LRRK2[R1441C] knock-in model), 6-month-old littermate matched were injected subcutaneously with vehicle [40% (w/v) (2-hydroxypropyl)-β-cyclodextrin (Sigma– Aldrich #332607)] or MLi-2 dissolved in vehicle at a 30 mg/kg final dose. Mice were killed by cervical dislocation 2 hours following treatment. Tissues were immediately harvested and snap-frozen in liquid nitrogen.

### Mouse genotyping

Genotyping of mice was performed by the MRC genotyping team at the MRC-PPU, University of Dundee, by PCR using genomic DNA isolated from ear biopsies. The VPS35[D620N] knock-in mouse strain required Primer 1 (5’ TCATTCTGTGGTTAGTTCAGTTGAG 3’), Primer 2 (5’ CCTCTAACAACCAAGAGGAACC 3’), and Primer 3 (5’ ATTGCATCGCATTGTCTGAG 3’) to distinguish wildtype from D620N knock-in alleles (60°C annealing temp). The LRRK2[R1441C] knock-in mouse strain required Primer 1 (5’ CTGCAGGCTACTAGATGGTCAAGGT 3’) and Primer 2 (5’ CTAGATAGGACCGAGTGTCGCAGAG 3’) to identify wildtype and R1441C knock-in alleles (60°C annealing temp). All genotyping primers were used at a final concentration of 10 pmol/μl. PCR reactions were set up and run using KOD Hot Start Polymerase standard protocol. PCR bands were visualized on Qiaexcel (Qiagen) using the standard DNA screening kit cartridge.

### Preparation of mouse tissue lysates

Snap frozen tissues were weighed and thawed on ice in a 10-fold volume excess of in icecold lysis buffer containing 50 mM Tris/HCl pH 7.4, 1 mM EGTA, 1 mM sodium orthovanadate, 10 mM 2-glycerophosphate, 50 mM NaF, 5 mM sodium pyrophosphate, 270 mM sucrose, supplemented with 1 μg/ml microcystin-LR, 1 mM sodium orthovanadate, complete EDTA-free protease inhibitor cocktail (Roche), and 1% (v/v) Triton X-100. Tissue was homogenized using a POLYTRON homogenizer (KINEMATICA), employing 3 rounds of 10 s homogenization with 10 s intervals on ice. Lysates were centrifuged at 20800 g for 30 min at 4°C and supernatant was collected for subsequent Bradford assay and immunoblot analysis.

### Quantitative immunoblot analysis

Clarified cell or tissue extracts were mixed with a quarter of a volume of 4× SDS–PAGE loading buffer [250 mM Tris–HCl, pH 6.8, 8% (w/v) SDS, 40% (v/v) glycerol, 0.02% (w/v) Bromophenol Blue and 4% (v/v) 2-mercaptoethanol]. 10-30 μg of samples were loaded onto NuPAGE 4–12% Bis–Tris Midi Gel (Thermo Fisher Scientific, Cat# WG1403BOX) and electrophoresed at 130 V for 2 h with the NuPAGE MOPS SDS running buffer (Thermo Fisher Scientific, Cat# NP0001-02). At the end of electrophoresis, proteins were electrophoretically transferred onto the nitrocellulose membrane (GE Healthcare, Amersham Protran Supported 0.45 μm NC) at 100 V for 90 min on ice in the transfer buffer (48 mM Tris–HCl and 39 mM glycine). Transferred membrane was blocked with 5% (w/v) skim milk powder dissolved in TBS-T [20 mM Tris–HCl, pH 7.5, 150 mM NaCl and 0.1% (v/v) Tween 20] at room temperature for 1 h. The membrane was typically cropped into three pieces, namely the ‘top piece’ (from the top of the membrane to 75 kDa), the ‘middle piece’ (between 75 and 30 kDa) and the ‘bottom piece’ (from 30 kDa to the bottom of the membrane). The top piece was incubated with rabbit anti-LRRK2 pS935 UDD2 antibody multiplexed with mouse anti-LRRK2 C-terminus total antibody diluted in 5% (w/v) skim milk powder in TBS-T to a final concentration of 1 μg/ml for each of the antibody. The middle piece was incubated with mouse anti-GAPDH antibody diluted in 5% (w/v) skim milk powder in TBS-T to a final concentration of 50 ng/ml. The bottom pieces were incubated with rabbit MJFF-pRab10-clone-1 monoclonal antibody multiplexed with mouse MJFF-total Rab10clone-1 monoclonal antibody diluted in 2% (w/v) bovine serum albumin in TBS-T to a final concentration of 0.5 μg/ml for each of the antibody. All blots were incubated in primary antibody overnight at 4°C. Prior to secondary antibody incubation, membranes were washed three times with TBS-T for 10 min each. The top and bottom pieces were incubated with goat anti-mouse IRDye 680LT (#926-68020) secondary antibody multiplexed with goat anti-rabbit IRDye 800CW ((#926-32211) secondary antibody diluted in TBS-T (1:10, 000 dilution) for 1 h at room temperature. The middle piece was incubated with goat anti-mouse IRDye 800CW (#926-32210) secondary antibody diluted in TBS-T (1: 10, 000 dilution) at room temperature for 1 h. Membranes were washed with TBS-T for three times with a 10 min incubation for each wash. Protein bands were acquired via near infrared fluorescent detection using the Odyssey CLx imaging system and quantified using the Image Studio software.

### Limit of detection experiments

The generation of A549 Rab10 [18] and Rab8A [7] knock-out cells have been described previously (S2 A). 10 mg of wild type, Rab8A knock-out and Rab10 knock-out cell extract was reduced by adding final 10 mM DTT (Dithiothreitol) and incubated 56 °C for 20 min and alkylated by adding 20 mM Iodoacetamide. After incubating in the dark for 30 min at room temperature proteins were acetone precipitated, resolubilized in 6M Urea buffer in 50 mM TEABC pH 8.0, digested using trypsin (1:20 ratio) and desalted using C18 cartridges as described previously [32]. The peptide amounts were determined by 3-(4-carboxybenzoyl)quinoline-2-carboxaldehyde (CBQCA) method [33]. 150 μg aliquots of each digest were lyophilized and stored at −80 °C.

### Bovine Serum Albumin tryptic digest

30 mg of Bovine serum Albumin was dissolved in 6 M Urea in 50 mM TEABC pH 8.0 buffer, reduced and alkylated using 5 mM DTT and 20 mM Iodoacetamide followed by incubation 30 min in the dark. The Urea concentration was reduced to 2 M using 50 mM TEABC pH 8.0 and digested with trypsin (1:20 ratio) at room temperature for 16 h. The digest was acidified by adding TFA to 1 % (v/v) and desalted using C18 Sep-Pak cartridges ([32]. Peptide amounts were determined by CBQCA assay [33] and 2 mg aliquots were lyophilized and stored at −80 °C.

### Pan-pRab immunoprecipitation to assess Rab1 phosphorylation by immunoblotting

To assess Rab1 phosphorylation, 12.5 μg of the Pan-pRab antibody was coupled to 25μL of Protein A/G Sepharose beads for 1 hour at 4°C with agitation. This was incubated with 2.3 mg of indicated MEF extract for 2 hours at 4°C. The immunoprecipitate was washed three times with lysis buffer. The immunoprecipitae was incubated in 20 μL of 2X NuPAGE LDS sample buffer and centrifuged through Spin-X column to remove resin. The eluate was used for detecting Rab1-Total (80% of sample) and Rab10-Total (20% of sample) by immunoblotting. We also undertook validation of novel stressMarq biosciences Rab1 total antibody (Fig S5 C). Lysates of HEK293 cells overexpressing HA-Rab1A, HA-Rab1B, HA-Rab1C or HA-empty were used for HA immunoprecipitation (300 μg per IP). Immunoprecipitates were then immunoblotted with the Rab1-total and ant-HA antibodies. Immunoblots were subjected to quantitative LI-COR immunoblot analysis with all indicated antibodies at 1 μg/mL. The Rab1 antibody detects Rab1A and Rab1B equally and Rab1C significantly more weakly (Fig S5 C).

### S-Trap assisted on-column digestion

Cell or tissue lysates (typically 250 μg) prepared as described above, was reduced by adding 10 mM Tris(2-carboxyethyl)phosphine(TCEP) and incubated at 60°C for about 30 min on a Thermomixer with a gentle agitation (800rpm). The samples were allowed to cool to room temperature and alkylated by adding 40 mM Iodoacetamide and incubated in dark at room temperature for 30 min on a Thermomixer by gentle agitation (800 rpm). The samples were supplemented sodium dodecyl sulfate (SDS) to 5% (v/v) from a stock solution of 10% (v/v) SDS and acidified by adding phosphoric acid to 1.2% (v/v) using 12% (v/v) phosphoric acid stock. The samples were immediately added to S-Trap wash buffer (90% (v/v) methanol in 100 mM TEABC pH 7.1) was added (7 times the volume of lysate). Under these conditions proteins form fine suspension particles and the resulting mixture was loaded on S-Trap mini columns. The columns were then centrifuged for 1 min at 1000 g, washed four times with STrap Wash buffer and transferred to 2 ml collection tubes. An “in-solution on-column” tryptic digestion was performed by applying 160 μl of 50 mM TEABC pH 8.0 containing a 1:15 trypsin: protein substrate applied to S-Trap column. The columns were spun down at 300g 1 min and the flow through reapplied to the columns and incubated on a Thermomixer at 47°C for 80 min followed by 3 h at room temperature without agitation. The columns were transferred to a new 1.5 ml low binding Eppendorf tube for the elution. 80 μl of 50 mM TEABC pH 8.0 was added and spun down at 1000g for 1 min. 100 μl of 0.15% (v/v) Formic acid was added to the columns and the spun down at 1000g 1 min. Another 100 μl of elution buffer 3 [80% (v/v) Acetonitrile and 0.15% (v/v)] was added and spun down at 500g for 3 min to completely elute the tryptic peptides. To ensure complete recovery, the columns were transferred to the previous collection tubes and another 100 μl of this elution buffer 3 was added and spun down. The eluates were combined, and peptides quantified using the CBQCA assay. For the total Rab protein analysis 0.5 μg of the tryptic digest aliquoted. For the phospho-Rab protein analysis 150 μg of tryptic peptide analysis was aliquoted for all cells and tissues other than brain in which 600-650 μg of tryptic peptide was used. Samples for total Rab protein analysis and the phospho-Rab analysis were vacuum dried in a speed-vac and stored in −80 °C until MS analysis.

### Preparation of Heavy stable isotope peptides

All heavy and light stable isotope synthetic peptides described in supplementary table 1 were synthesized by JPT peptide technologies (https://www.jpt.com/) in 1 nano mole aliquots. All the synthetic peptides were quantified by amino acid analysis and LC-MS analysis by JPT and confirmed to be of purity of >95%. The peptides were delivered in a lyophilized form and were resuspended in solvent containing 0.1% (v/v) formic acid in 3% (v/v) acetonitrile to give a final concentration of 10 pmol/μl. Aliquots of this were further diluted in a series of 10-fold dilution to a lowest concentration of 10 fmol/μl and stocks of each dilution aliquoted and stored at −80 °C.

### Immunoprecipitation of pRab, total LRRK2, pSer910 and pSer935 peptides

Aliquots of vacuum dried peptides (0.5 μg for total Rab samples and 150 μg for phospho-Rab samples) were dissolved in 1 ml of Immunoaffinity purification buffer (50 mM MOPS, 50 mM Na2HPO4 and 150 mM NaCl). The samples were sonicated for 10 min at room temperature, centrifuged at 20,800g for 10 min and transferred to a 1.5 ml low binding Eppendorf tube. A 10 μl mixture containing 100 fmol of each of the pRab1, pRab3, pRab8, pRab10, pRab35, pRab43, tLRRK2(388-399), LRRK2-s910 and LRRK2-s935 heavy isotope synthetic peptide mixture was added. Next an 10 μl suspension of Immunoaffinity purification buffer containing 50% (v/v) Protein A/G resin conjugated to 0.5 μg PAN-pThr-Rab antibody (pRab8), 0.5 μg LRRK2 total antibody (8G10) antibody, 0.5 μg LRRK2-pS910 (UDD1) and 0.5 μg LRRK2-pS935 (UDD2) was added. Samples were incubated on an end-to-end rotator in 4° C for 2h, centrifuged at 2000g for 2 min at 4 °C. The supernatant was collected into new 1.5 ml Eppendorf tubes and subjected to a second round of immunoprecipitation as described above. The immunoprecipitates washed sequentially 1 ml of Immunoaffinity purification buffer, 2 washes with 1 ml of phosphate-buffered saline, and finally with 1 ml of Milli-Q water and the supernatant removed to leave just the immunoprecipitate. 60 μl of 0.2% (v/v) TFA was added to the washed immunoprecipitates and the resulting mixture incubated at room temperature on a Thermomixer at 1200 rpm for 10 min to elute absorbed peptides. The tubes were centrifuged at 2000 g for 2 min and the supernatant containing the eluted peptides collected in a new 1.5 ml Eppendorf tube. The resin was washed again with 40 μl of 0.2% (v/v) TFA and the supernatants containing the immunoprecipitated peptides were combined.

### Preparation of samples for total Rab peptide quantitation

0.5 μg of peptide digest was dissolved in 60μl of buffer containing (0.1% (v/v) formic acid in 3%(v/v) acetonitrile and sonicated at room temperature for 10 min. Further the peptide digest spiked with an equimolar ratio of 50 fmol of heavy labelled peptide mixture containing Rab1A, Rab1B, Rab3D, Rab8A, Rab8B, Rab10, Rab12, Rab35 and Rab43. The sample was then loaded on EvoTips as described below were either subjected to LC-MS/MS analysis immediately or stored at 4°C.

### C18-Peptide clean-up of samples

C18 clean-up of the immunoprecipitated peptides prior to MS analysis employing in-house prepared Stage tips. A single 18-gauge syringe was used to punch a hole in C18 material from 47 mm think C18 disk which was packed into a 250 μl pipette tips. The following solvents were prepared fresh, Activation solvent (100% (v/v) Acetonitrile); Solvent-A (0.1% (v/v) TFA); Solvent-B (50% (v/v) Acetonitrile in 0.1% (v/v) TFA). 60 μl of Activation solvent was added to the stage tip which was centrifuged at 2000g for 2 min. Flow through was discarded and the stage-tips were equilibrated by adding 60 μl of Solvent-A followed by centrifugation at 2000g for 2 min and this step was repeated a further time. The acidified peptides were loaded onto the stage tip, which was centrifuged at 1500g for 4 min. The flow through was reapplied and centrifuged through the tip a second time to ensure all peptides were absorbed onto the C18 resin. The stage-tips were next washed twice by centrifugation of 60 μl of Solvent-A through the tip. Finally, the stage-tips were transferred into a new collection tubes and the peptides were eluted by adding 30 μl of solvent-B and centrifuged and this step was repeated once again, and the eluates combined. The eluate was flash frozen on a dry-ice and vacuum dried and stored in −80 °C freezer until the LC-MS/MS analysis.

#### Deep proteomic profiling of wild type MEFS

300 μg of wild type MEFs were prepared for tryptic digestion using S-Trap assisted on-column digestion workflow as described above. Eluted peptides were vacuum dried and subjected to basic reverse-Phased liquid chromatography fractionation as explained previously [32]. A total of 96 fractions were collected and concatenated to 48 fractions and further vacuum dried and stored in −80 °C deep freezer until further LC-MS/MS analysis.

### Preparation of Evotips

The vacuum dried peptides were resuspended in 60 μl of Solvent (0.1% (v/v) Formic acid in 3% (v/v) Acetonitrile) and sonicated on a water bath sonication for 4 min at room temperature. In addition, XX μl of a 25 Fmol of Pierce retention time correction (PRTC, Thermo Fisher Scientific #88320) was spiked-in to the mixture to correct on-the-fly scheduled retention times of the targeted analytes on the mass spectrometer. Evotips were purchased from EvoSep (Odense Denmark #EV2001). The following solvents were prepared in fresh. Solvent-A (0.1% (v/v) Formic acid); Solvent-B (100% (v/v) Acetonitrile). The Evotips were activated by adding 20 μl of Solvent-B which was centrifuged through the tip at 1000g for 1 min. The Evotips were next immersed in a 200 μl of Isopropanol for few seconds to eliminate any polymers. The tips were next equilibrated by adding 20 μl of Solvent-A which was centrifuged through at 1000 g for 1 min and this step repeated once more and the eluates discarded. The acidified peptides for the total Rabs and immunoprecipitation samples were loaded onto the Evotips which were next centrifuged at 700 g for 2 min. The Flow through was reapplied to the tip and centrifuged again for 700 g for 2 min. The Evotips were washed by adding 20 μl of Solvent-A, centrifuged at 1000g for 1 min and this step repeated a further time. A final 100 μl of Solvent-A was added and centrifuged for 4 seconds such that the Solvent-A protects the C18 material from evaporation during the LC-MS/MS analysis. At this stage, the prepared EvoTips are transferred onto the 96 well plate box and placed on EvoSep auto sampler tray.

### LC-MS/MS analysis of immunoprecipitated peptides

Unless stated otherwise PRM data was acquired with the following settings. EvoTips were loaded on EvoSep LC system auto sampler tray and analyzed using 21 min (60 samples per day) script. The EvoSep LC system elutes the peptides from EvoTip using low pressure pump-A and pump-B by directly applying 35% solvent B (Acetonitrile in 0.1% (v/v) formic acid) the eluted peptides stored in a long capillary loop where using Pump-C and Pump-D generates a pre-formed gradient that applies the peptides into the analytical column Reprsoil-pur C18 AQ, 1.5um beads, 150um ID, 8cm long for pRab analysis(21 min script) and Reprsoil-pur C18 AQ, 1.9um beads, 150um ID, 15cm long for total Rab analysis (44min script). The eluted peptides are sprayed into the mass spectrometer that was operated in a PRM mode using Xcalibur software interface by including one Full MS scan within 300-800 m/z per duty cycle. The Full MS acquired at 120,000 resolution at m/z 200 by accumulating 3E6 ions in the C-trap for a maximum of 20 ms. The targeted MS2 (PRM) scans were performed by importing inclusion list containing pRabs, total and phospho LRRK2 and PRC peptide sequences. The inclusion list containing scheduled retention times, +2 or +3 charge states are provided in supplemental Table 2. The PRM MS2 scans for each targeted ion was isolated using 0.7 Da Quadrupole filter and 1E5 AGC target was set and accumulated in the C-trap for 250 ms by using 27% higher energy collisional dissociation (HCD). The MS2 scans were acquired at 30,000 resolution at 200 m/z analyzed using Ultra-high field Orbitrap mass analyzer. The loop count was set to 12 scans per duty cycle. Dynamic retention time algorithm was enabled to correct on-the-fly scheduled retention times of the target ions.

The retention times of pRabs were determined by using an equimolar mixture of 50 fmol of pRab1, pRab3, pRab5, pRab8, pRab12, pRab29 Thr71, pRab29 Ser72, pRab29 Thr71+Ser72, pRab35 and pRab43 peptide a loaded on Evotips as described above. The samples were analyzed in a data dependent acquisition mode (DDA) by disabling dynamic exclusion option on a QE HF-X instrument that is in line with the EvoSep LC system. Full MS was acquired at 60,000 resolution (m/z) and measured in an Orbitrap mass analyzer. MS2 scans were performed by selecting top 10 precursor ions were isolated at 0.7 m/z Isolation width using segmented Quadrupole and subjected to normalized higher energy collisional dissociation of 27% (HCD) and measured using Orbitrap mass analyzer at 30,000 resolution (m/z 200). The AGC targets, 3E6 and 5E4 and Ion filling times 20 ms and 40 ms were kept for MS1 and MS2 respectively.

### LC-MS/MS analysis for total Rab peptides

For total Rab peptides samples were analyzed as above with the exception that peptides were separated on a 15cm analytical column by employing 44 min script on EvoSep LC system. The Mass spec data was acquired in a PRM mode as explained above. The retention times of total Rab peptides were determined by using an equimolar mixture of 50 fmol of optimized Rab1A, Rab3D, Rab8A, Rab8B, Rab10, Rab12, Rab35 and Rab43 peptides (Supplementary Table 2)

### LC-MS/MS analysis limit of detection studies

For the Limit of detection of Rab10 and Rab8A knock-out A549 cell extract and HeLa cell extract the samples were prepared as explained above by employing 21 min run script on EvoSep LC system. Only the heavy and light pRab10 Thr73 (FHp**T**ITTSYYR, for z=2 m/z 684.8028, m/z 689.8070, and z=3 m/z 456.8710, m/z 460.2071) and only the heavy and light pRab8 Thr72 (FRp**T**ITTAYYR for z=2 m/z 686.3265, m/z 691.3306 and z=3 m/z 457.8867, m/z 461.2228) with a scheduled retention times of ± 4 min window was specified. The Mass spectrometers was operated in a targeted MS2 (PRM) mode by including single Full MS and the data was acquired as described above. MS2 scans were acquired by specifying an AGC target of 1E5 ions accumulated either for 500 ms or 300 ms by applying 27% NCE of HCD and measured using Ultra high-field Orbitrap mass analyzer at 30,000 resolution (m/z 200).

### LC-MS/MS analysis for higher-energy collisional dissociation (HCD) studies

Samples were prepared as described above by employing by 8 cm column and 21 min run script on EvoSep LC system and Full MS and targeted MS2 (PRM) parameters were used except the HCD was varied at 25%, 27% and 30% energies.

### Deep proteomic profiling of wild type MEFs studies

48 bRPLC fractions were prepared for LC-MS/MS analysis by dissolving each fraction in 60 μl of 0.1% formic acid (v/v)in 3% (v/v) aceotonitrile buffer and sonicated for 10 min at room temperature. 15 μl of each fraction was analyzed on Orbitrap Exploris 480 mass spectrometer that is in-line with Dionex 3000 RSLC nano liquid chromatography system. Peptides were loaded on to 2cm trap column (C18, 5 μm, 100 Å, 100 μm × 2 cm; PN: 164562, Thermo Scientific) and further loaded on 50 cm Easynano spray analytical column (2 μm, 100 Å, 75 μm × 50 cm. PN: ES803, Thermo Scientific) and electro sprayed directly into the Mass spectrometer using Easynano source. Peptides were separated by employing a gradient containing buffer A (0.1% Formic acid (v/v) and buffer B (80% (v/v) acetonitrile in 0.1% formic acid (v/v) by employing a gradient consists of 7% B to 40% B for 70 min and further the column was kept at 90% B for 10 min and equilibrated with 3% B for additional 20 min. The mass spectrometer was operated in a data dependent mode in a top speed mode for 2 seconds. Full MS was acquired at 60,000 resolution at m/z 200 and measured using Orbitrap mass analyzer. The peptides were isolated by using 1.2 Da isolation width using Quadrupole mass filter and fragmented using 30% normalized HCD. MS/MS was acquired at 15,000 resolution at m/z 200 and measured using Orbitrap mass analyzer. AGC target for MS was kept at 300 % =3E6 ions and for MS/MS 100% = 1E5 ions by accumulating for a maximum of 20 ms for MS and 55 ms for MS/MS.

### LC-MS/MS analysis for antibody validation

Samples were prepared as described above by employing a 15cm column with a 44 min or 60 min run in a DDA mode on QE-HFX mass spectrometer under the control of MaxQuant live application [34]. Full MS was acquired at 120,000 resolution at (m/z200) by specifying 1E6 AGC target and a maximum ion injection times of 20 ms. Top 10 precursors were selected from the survey scan and were fragmented using 27% HCD and measured at 15000 resolution (m/z 200) by specifying 1E5 AGC target for 28 ms ion injection times.

### Spectral library generation

Raw data was processed using Proteome Discoverer version 2.2 software and searched using Sequest algorithm against Human/Mouse Uniprot databases (Uniprot release 2017_03) that were appended with Pierce retention time calibration mix peptide sequences. The data was searched with a 10 ppm mass tolerance of MS1 and 0.05 Da MS2 tolerance by allowing 2 missed cleavages. Protease was specified as trypsin with a minimum 7 Amino acid sequence length. Oxidation of Met, deamidation of Asn/Gln, Phosphorylation of Ser/Thr, Heavy lysine (^13^C_6_ ^15^N_2_), Heavy Arginine (^13^C_6_ ^15^N_4_) were set as dynamic modifications and Carbamidomethylation of Cys was set as fixed modifications. 1% PSM, peptide and protein FDR was applied by validating with percolator node. Either the Proteome Discoverer result or Thermo .msf files were directly imported into Skyline to build the spectral libraries for indicated pRabs, total Rab’s, total LRRK2 and phospho LRRK2 and PRTC peptide sequences.

### Database searches

Raw data was processed using MaxQuant (version 1.6.0.13). The data was searched against either Uniprot (Human or Mouse 2017_03 build) using in-built Andromeda search engine. Following group specific parameters were used, Trypsin as a protease, Oxidation of Met, Acetyl-Protein N-terminal, Deamidation of Asn/Gln, Phosphorylation Ser/Thr residues were selected as dynamic modifications. Carbamidomethylation of Cys as a fixed modification. LFQ min ratio count was set to 1. 20 ppm of First search tolerance and 4.5 ppm of main search tolerance was applied. 1% PSM and protein FDR was applied. MaxQuant search output tables were processed using Perseus software suite for downstream Bioinformatic analysis. Protein copies were estimated by using Proteomic ruler plug-in within Perseus software suite.

### Data analysis

The multiplexed pRabs and total LRRK2 targeted PRM assay falls into tier-2 assay as per the previously published guidelines [35, 36]. Spectral libraries were generated as described above which were used to identify and to quantify the top six fragment ions. Mass spectrometry raw data was directly imported into Skyline software suite (version: 19.1.0.193). The target peptides were manually entered by specifying the corresponding gene names. Skyline software automatically picks the chromatographic traces and peak boundaries. The graphical chromatographic traces, peak boundaries and retention time alignments for every target ion were further manually verified and further any interfering y1 or b2 ions were excluded. Samples that showed variation in Internal standard due to the losses originated in immunoprecipitation or C18 stage-tip clean up were excluded and indicated in the corresponding supplemental table. Synchronization of isotope label types were enabled such that any transition that was less specific could be removed in both light and heavy labelled fragment ions. Precursor charge states of 2,3, ion charges 1, 2 and ion types of y,b,p were selected in peptide transition settings. Quantification was performed in determining the relative abundance was performed on both MS1 and MS2 levels. Ratio to the stable isotope internal standard values were exported into the excel sheet to determine the relative abundance of each target peptide. The estimated phospho Rab and total LRRK2 and phospho LRRK2 abundance was calculated as described previously [37], (Light/Heavy ratio multiplied to the Spike-in amount). The relative fold expression values were determined by Light/heavy ratio values between pathogenic LRRK2 or VPS35[D620N] mutants versus the corresponding wild type samples groups. For total Rab PRM analysis, peak integration was carried out using Skyline software and extracted ion chromatograms were manually verified for and any interfering ions of y1 or b2 ions were excluded. Light peptide intensities were exported into excel sheet for further analysis. For Rab proteins where >1 unique peptide was monitored the summed intensities were used. Relative levels of each Rab protein was determined by dividing the peptide intensity to the median intensity of the corresponding peptide across conditions and replicates. All of the quantitation and summary of the results provided in supplementary table 2 and 3.

Bioinformatic and statistical analyses was carried out in this study using Microsoft excel, Perseus software suite and GraphPad prism (version 8.4.3) which includes the quantification results of both Immunoblotting and Mass spectrometry analyses. Finally all figures are organized in Adobe illustrator CC version 21.1.0.

## Results

### Relative abundance of LRRK2 pathway components in mouse embryonic fibroblasts

We first generated a deep proteomic profile of ~10,000 proteins in mouse embryonic fibroblasts that are widely used to study LRRK2 signalling using high resolution MS (Fig 1A, B, Table S2). Employing the histone-based proteomic ruler approach [38], we estimated the copy number of the known, key LRRK2-Rab pathway components (Fig 1A,B). From highest to lowest abundance we detected Rab1A and Rab10 (~1.25 million copies per cell), Rab8A and Rab1B (~0.65 million copies per cells), Rab8b, Rab12 and Rab35 (~0.2 million copies per cells), Rab3A, Rab3D, Rab43 and Rab3B (~70,000 copies per cell), Rab29 (20,000 copies per cell). LRRK2 was lower abundance (6000 copies per cell) and ranked at 7320 on our list of 10,000 most abundant proteins. Other LRRK2 pathway components included VPS35 (~0.7 million copies per cell), RILPL1 (~56,000 copies per cell), RILPL2 (~5000 copies per cell), JIP3 (~2,000 copies per cell), JIP4 (~0.13 million copies per cell) and PPM1H (~10,000 copies per cells). LRRK1 was expressed at lower levels than LRRK2 (~3,000 copies per cell).

**Figure 1:**
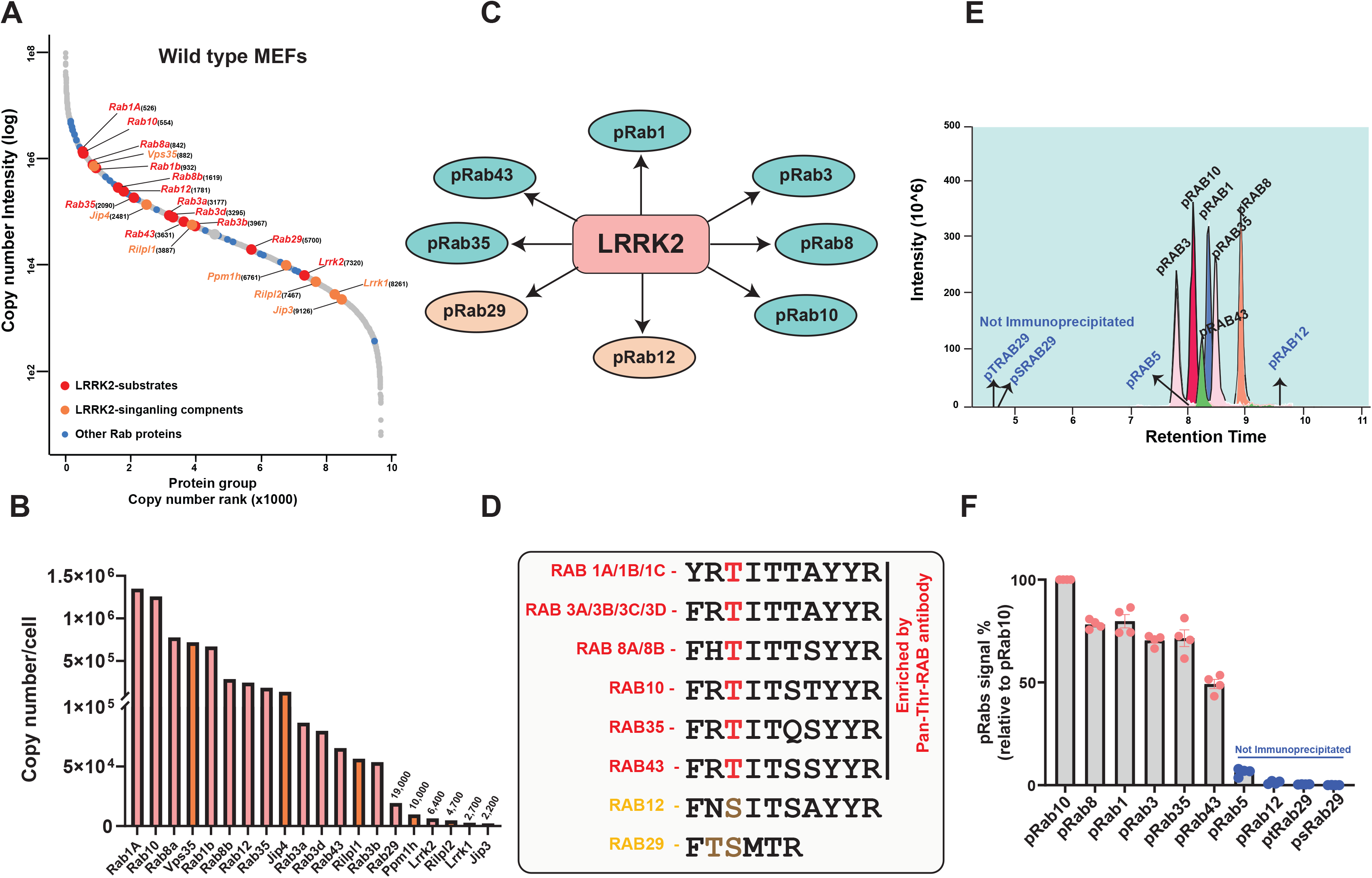
Identification and quantification of endogenous LRRK2 signalling components in mouse embryonic fibroblasts and development of a targeted MS assay to enrich pRab peptides. **A)** 250 μg of wild type MEFs extract was tryptic digested, high pH fractionated and analysed on an Orbitrap Exploris mass spectrometer. Protein copy numbers per cell were estimated using the histone-based Proteomic ruler. The relative copy number rank is depicted in the rank abundance plot. LRRK2-phosphorylated Rab proteins are indicated in red coloured text along with their corresponding rank number in parentheses. Other LRRK2 signalling components are indicated in orange coloured dots and texts. **B)** Relative copies of LRRK2 signalling components depicted in bar chart, y-axis indicating the protein copies per cell for each of the indicated protein group on x-axis. **C)** Endogenous Rab substrates that are phosphorylated by LRRK2. **D)** Sequence alignment of LRRK2-phosphorylated switch-II Thr and Ser Rab proteins. Rab proteins that possess switch-II Thr phosphorylation site (Rab1, Rab3, Rab8, Rab10, Rab35 and Rab43) that are immunoprecipitates by Pan-pRab antibody are highlighted in red coloured text. Rab29 (Thr/Ser) and Rab12 (Ser) proteins are highlighted in orange coloured text. **E)** 100 fmol of equimolar heavy stable synthetic phosphoRab peptide mixture (pRab1, pRab3, pRab5, pRab8, pRab10, pRab12, pRab29 (Thr), pRab29 (Ser), pRab (Thr/Ser) and pRab43), spiked into 200 μg of bovine serum albumin tryptic digest and was subjected to peptide immunoprecipitation using Pan-pRab antibody. Data was acquired on Orbitrap HF–X MS in a targeted PRM mode. Extracted ion chromatograms (XIC) depicting the relative abundance of each of the Rab Phosphopeptide. X-axis indicating the retention time elution of six pRabs (Rab1, Rab3, Rab8, Rab10, Rab35, pRab43) eluting between 7.5 to 9 min of a 21 min run on EvoSep LC-system. pRab peptides that are not immunoprecipitated by Pan-pRab antibody (pRab29(Thr), pRab29 (Ser), pRab5 (Ser) and pRab12 (Ser) are indicated in blue-coloured text. **F)**. Bar graph depicting the relative abundance of Figure E are depicted as a percentage of recovery relative to pRab10 signal. pRab peptides that are not immunoprecipitated by Pan-pRab antibody (pRab29(Thr), pRab29 (Ser), pRab5 (Ser) and pRab12 (Ser) are indicated in blue-coloured text. n=4, Error bars indicating the mean ± SEM.

### Development of a targeted MS assay to assess LRRK2-phosphorylated Rab proteins

As a first step towards developing a targeted MS assay to detect LRRK2-phosphorylated Rab proteins, we generated synthetic, heavy, stable isotope phospho-peptides corresponding to the tryptic peptide sequences of all reported LRRK2 substrate Switch-II phosphorylation sites (Table S1). These include peptides corresponding to Rab29 phosphorylated at Thr71 or Ser72 or phosphorylated at both Thr71 and Ser72 (Table S1). The amino acid sequences of the switch II motif peptides between each of the Rab protein is unique, but between isoforms including Rab 1(Rab1A/B), Rab3 (Rab3A/B/C/D), Rab(5A/B/C) and Rab8 (Rab8A/B) the sequences are identical and therefore cannot be distinguished (Fig 1C,D + Table S1). Furthermore, the monoisotopic mass of each of the pRab peptides is unique apart from Rab3 isoforms and Rab35 that are isobaric (Table S1). Mixtures containing 10 or 50 fmol of each phospho-peptide were subjected to C18 reverse phase chromatography on an EvoSep LC system employing a 21 min run [39]. Apart from the Rab29 peptides that elute between 1-2 min, too early to quantify, all the other pRab peptides were well separated, allowing MS identification and quantitation of each peptide on a QE-HFX instrument (Fig S1A). The isobaric pRab3 and pRab35 phospho-peptides elute over 1 min apart, enabling these sites to be discriminated (Fig S1A).

The retention times of the pRab peptides in independent chromatographic runs was highly reproducible in mixtures containing 10 fmol (Fig S1B) or 50 fmol (Fig S1C) pRab phosphopeptides. We compared the performance of two LC systems (the EvoSep and Dionex-RSLC3000) and found that the signal intensity (Fig S1D) and resolution of separation was superior on the EvoSep LC system. In addition, the carryover of samples between independent runs using the EvoSep was below 0.1% compared to 1-3% in the Dionex-RSlC3000 system (Fig S1E). The reduced carryover of the EvoSep system is due to utilization of a disposable tip-based trap column to load the sample. Moreover, the run to run overhead time between runs is 2 min on the EvoSep compared to 20 min on the Dionex-RSlC3000 system allowing significantly more samples to be analyzed per day using the EvoSep LC system.

### An immunoprecipitation enrichment step to isolate endogenous pRab tryptic peptides

As discussed earlier, due to the low stoichiometry of Rab phosphorylation an enrichment step prior to MS analysis is required to robustly quantify these peptides [18, 25]. Even for Rab10 that is the most abundant LRRK2-phosphorylated Rab protein, an SDS-polyacrylamide gel electrophoresis enrichment step was found necessary for accurate measurement of phosphorylation stoichiometry [25]. We therefore explored whether a previously described, commercially available phospho-specific antibody termed here “Pan-pRab antibody” could be exploited to enrich pRab peptides from tryptic digests of cell extracts. Previous work had revealed that this antibody immunoblotted and immunoprecipitated multiple endogenous Rab proteins phosphorylated on a Thr residue by LRRK2 [7, 26]. Test immunoprecipitations with an equimolar mixture of 100 fmol of the heavy labelled pRab peptides spiked into a 200 μg tryptic digest of bovine serum albumin, revealed that the pan-pRab antibody immunoprecipitated pRab1, pRab3, pRab8, pRab10 and pRab35 phospho-peptides quantitatively, whilst pRab43 was also immunoprecipitated with ~70% efficiency (Fig1E,F). Consistent with previous work [26], the immunoprecipitated pRab peptides all possess a Thr switch-II phosphorylation site. The pRab5, pRab12 and pRab29 peptides possessing a Ser Switch-II motif were not immunoprecipitated (Fig1E, F). Dose studies revealed that 0.1 to 1 μg of pan-pRab antibody was required to immunoprecipitate pRab1, pRab3, pRab8, pRab10 and pRab35 robustly (Fig S1F). Even at 1 μg of pan-pRab antibody, ~70% pRab43 was immunoprecipitated (Fig S1F).

We next analyzed the limit of detection of this immunoprecipitation approach by spiking in pRab8 synthetic heavy and light labelled peptides into a tryptic, 200 μg digest of an extract from Rab8A knock-out A549 cells [7] (Fig 2A, S2A). We also spiked equivalent pRab10 peptides into extracts of Rab10 knock-out A549 cells [18] (Fig 2B, S2A). For both extracts, the heavy labelled peptide was kept constant at 50 fmol whilst levels of the light peptide varied from 0.01 to 1000 fmol. The pRab peptides were immunoprecipitated using the pan-pRab antibody and analyzed as above following EvoSep chromatography and MS analysis. This revealed that for both pRab8 and pRab10 peptides, the limit of detection for high confidence quantitation is around 0.1 fmol (Fig 2A,B). We detected 0.01 fmol, but quantitation was outside a linear range. We also undertook the equivalent experiments adding synthetic heavy and light stable pRab8 and pRab10 peptides to 50 ng of HeLa cell tryptic digest which was analyzed without an immunoprecipitation step (Fig S2B, C). This analysis confirmed that 0.01 fmol of pRab peptide could be detected but for linear quantitation at least 0.1 fmol was required.

**Figure 2:**
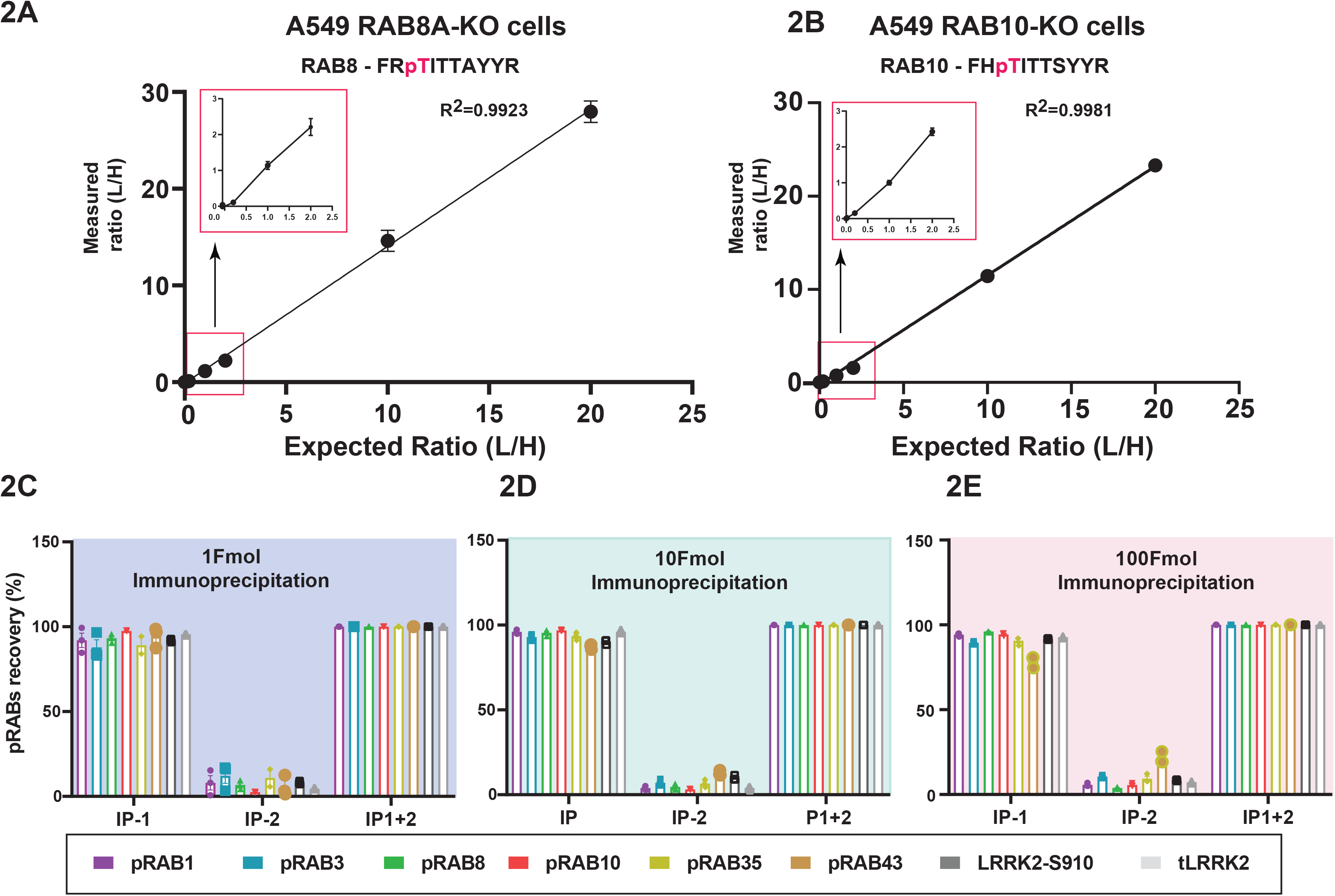
Development of a targeted MS assay to enrich endogenous phosphorylated Rab peptides. A) 250 μg of Rab8A-KO A549 cell extract was subjected to tryptic digestion. Limit of detection experiments were carried out by spiking synthetic light phosphorylated Rab8A (Thr72) peptide at (0.01, 0.1 1, 10, 50, 100, 500, 1000 fmol), while keeping the constant 50 fmol of heavy phosphorylated Rab8A (Thr72) peptide. Immunoprecipitations were undertaken with Pan-pRab antibody and analysed on Orbitrap HF-X MS in a targeted PRM mode. XIC were generated using Skyline software. X-axis depicting the expected (L/H) ratio and Y-axis depicting measured (L/H) ratio. Rectangular box indicates the zoom in of 0.01, 0.1 and 1 fmol data points. n=3. B) As in A) except that 250 μg of Rab10-KO A549 cell extract was utilized. C-E) 1, 10 and 100 fmol of heavy synthetic six Rab peptide mixture (pRab1, pRab3, pRab8, pRab10, pRab35, pRab43) and LRRK2-pS910 and total LRRK2 peptides were spiked into 200 μg of BSA tryptic digest and sequential immunoprecipitations(IP) were undertaken using Pan-pRab antibody. Levels of peptides eluted from the beads on the 1^st^ round (IP-1) and 2^nd^ round IP (IP-2) are indicated. Samples were analysed in a PRM mode. Data was analysed to assess the relative recovery of pRabs in each round of IP, N=3, error bars indicating the mean ±SEM.

### Expanding the assay to detect total and pSer910 and pSer935 phosphorylated LRRK2

We next explored whether the PRM MS assay could be expanded to quantify total LRRK2 as well as Ser910 (UDD1) and Ser935 (UDD2) phosphorylated LRRK2. Test immunoprecipitations were performed with three widely utilized LRRK2 total monoclonal antibodies as well as with anti-pSer910 and anti-pSer935 monoclonal antibodies. We used 200 μg of tryptic peptide digests derived from either mouse MEFs (Fig S3D) or human A549 cells (Fig S3A,D,E) that express relatively high levels of endogenous LRRK2. MS analysis of the immunoprecipitates revealed that one of the total LRRK2 antibodies (8G10) specifically immunoprecipitated high levels of a single tryptic peptide encompassing residues 390-401 of mouse LRRK2 (IGDEDGQFPAHQ) and the equivalent human peptide (388-399, IGDEDGHFPAHR) (Fig S3A). Further, independent epitope mapping analysis confirmed that the 8G10 antibody detected the peptide sequence DEDGHFP (Fig S3B). The other total LRRK2 antibodies tested immunoprecipitated lower levels of several peptides containing missed-tryptic cleavages that are not ideal for a targeted assay (FigS3A). The anti-pSer910 (UDD1) and anti-pSer935 (UDD2) monoclonal antibodies also immunoprecipitated a single LRRK2 tryptic peptide encompassing the expected phosphorylated residues in both mouse and human cells (Fig S3-A,D,E). The peptides immunoprecipitated with the 8G10 and pSer910 antibodies possess a single amino acid difference and anti-pSer935 peptide possesses a two amino acid conservative change between the human and mouse sequences (Fig S3C). We therefore generated synthetic heavy stable isotope peptides encompassing the mouse and human sequences of the tryptic peptides targeted by the 8G10, pSer910 and pSer935 LRRK2 monoclonal antibodies (Table S1). These were analyzed on an EvoSep LC system in a pool in combination with an equimolar amount of the pRab peptide mixture. The peptides were all well separated in a 21 min run, allowing MS identification and quantitation of each peptide.

To optimize detection further, we compared 3 different collision energies (HCD25, HCD27 and HCD30) employing 10 fmol and 50 fmol of each peptide in 50 ng of HeLa cell tryptic peptide digest (Fig S4A-D). For each peptide a coefficient of variation was measured in triplicate experiments. HCD27 was observed to provide the optimal collision energy to assess this set of peptides. It should be noted that Rab43 had the highest variation (~12%) while the Rab peptides displayed a median coefficient of variation of 4.5% (Fig S4A to D). All subsequent analysis undertaken in this study was therefore performed at HCD27.

### Multiplexing of pan-pRab, 8G10, pSer910, pSer935 and antibodies

We next multiplexed the pan-pRab, 8G10, pSer910, pSer935 and antibodies (0.5 μg each) to verify whether these could efficiently immunoprecipitate the pRab peptides using mouse (Fig 2C,D,E) or human (Fig S2D,E,F) peptides. Test immunoprecipitations were undertaken with an equimolar mixture of 1 fmol (Fig 2C, S2D), 10 fmol (Fig 2D, S2E) and 100 fmol (Fig 2E, S2F) of each of the peptides spiked into a 200 μg tryptic digest of bovine serum albumin. To maximize recovery of Rab43, we compared recovery of each peptide after 2 rounds of immunoprecipitation. This analysis revealed that all peptides in the 1-100 fmol tested conditions, including Rab43, were efficiently immunoprecipitated by >95% after 2 rounds of immunoprecipitation; we could also readily detect and quantify mouse (Fig 2C,D,E) and human (Fig S2D,E,F) peptides.

### Quantitation of total Rab proteins in cell extracts

As Rab proteins are much more abundant than LRRK2 (Fig 1A,B), we next explored whether we could directly detect and quantify signature tryptic peptides derived from individual Rab proteins straight in tryptic digests of 500 ng of wild type MEFs analyzed using the EvoSep LC system on the QE-HFX instrument. Using a 45 min run, we reliably detected tryptic peptides derived from Rab1A, Rab1B, Rab3D, Rab8A, Rab8B, Rab10, Rab12, Rab35 and Rab43 at a high signal to noise range; Rab29 and LRRK2 that are expressed at low levels (Fig 1A,B) were below the detection limit. In a 21 min run, levels of Rab8B, Rab35 and Rab43 that are less abundant than the other Rabs (Fig 1A,B), were harder to detect and quantify reliably. We therefore adopted the 45 min run protocol. For each Rab protein, we selected between 1 and 3 unique proteotypic peptides of the same sequence in mouse and human that based on our proteome analysis, permits robust detection and quantitation of Rab1A, Rab1B, Rab3D, Rab8A, Rab8B, Rab10, Rab12, Rab35 and Rab43 proteins. The sequence and m/z of these peptides are listed in S. Table1. We synthesized heavy stable isotope peptides for each of these Rab proteins and spiked 50 fmol (Fig S5A) or 10 fmol (Fig S5B) of these peptide mixtures into 50 ng of Hela tryptic digest. These peptides separate well on a 45 min EvoSep run, allowing quantitation of Rab proteins in the tryptic digest without further enrichment (Fig S5A,B).

### Workflow to quantify phosphorylated and total LRRK2 and Rab proteins in cell and tissue extracts

Based on the groundwork described above, we developed a workflow to assess endogenous levels of total and LRRK2-phosphorylated Rab1, Rab3, Rab8, Rab10, Rab35 and Rab43 as well as total LRRK2 and LRRK2-phosphorylated Ser910 and Ser935 (Fig 3). In this workflow, cells or tissues are lysed with conventional lysis buffer containing detergents such as 1% (v/v) Triton-X100 required to solubilize Rab proteins in addition to phosphatase inhibitors. For MEFs, mouse liver, kidney and spleen extracts that express relatively high levels of LRRK2, we typically used 0.15-0.3 mg per sample. For mouse brain that expresses lower levels of LRRK2, we used 1 mg extract per sample.

**Figure 3:**
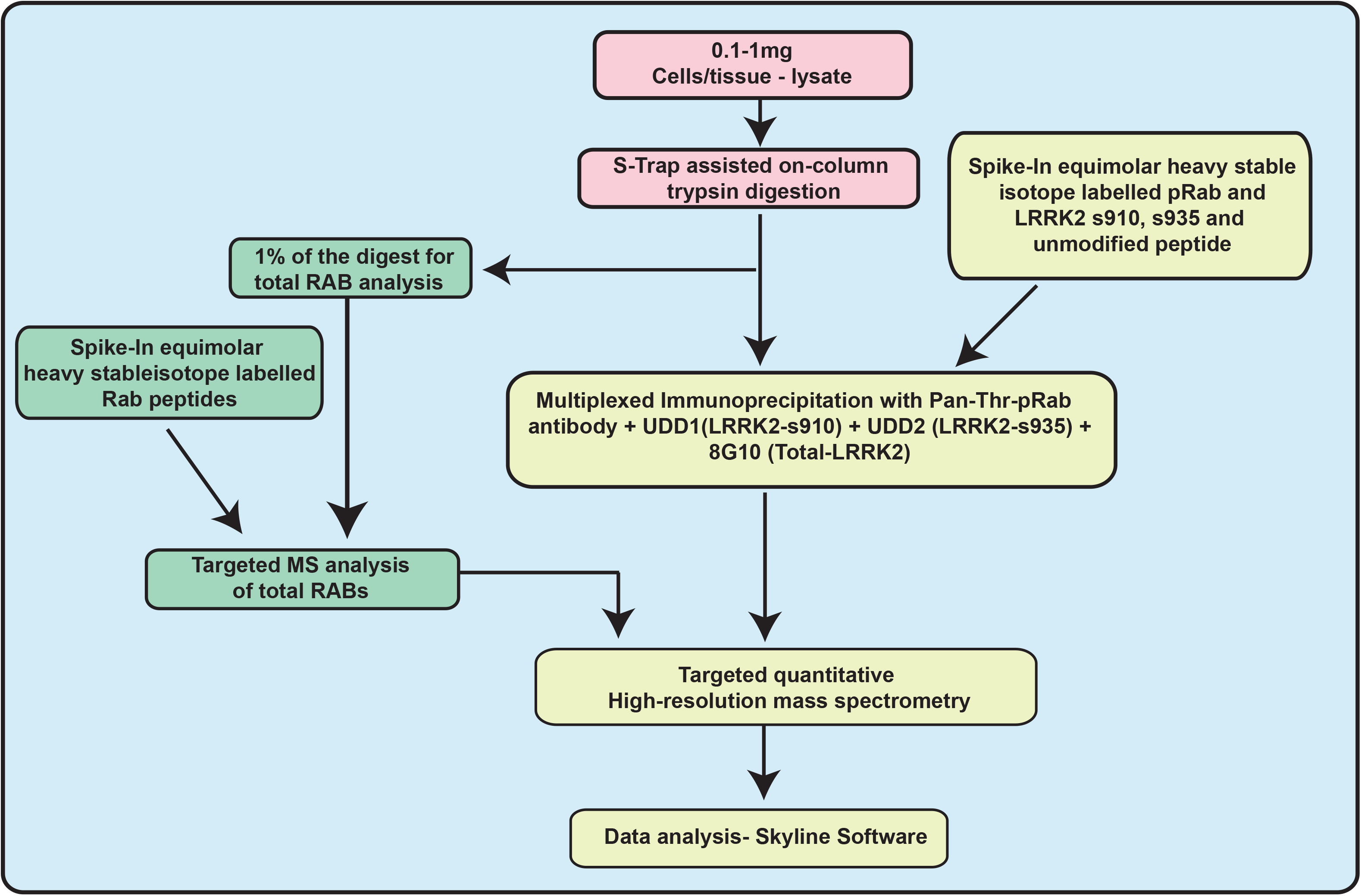
Workflow of targeted LRRK2-Rab Pathway assay. Summary of workflow we have developed to quantify endogenous levels of pRab, total Rab, LRRK2-pS910/pS935 and total LRRK2 peptides in cells and tissue extracts.

The cell/tissue extract is reduced, alkylated and absorbed onto an to S-Trap (suspension trap) spin column [40], which is washed to remove detergents and non-protein debris. Tryptic digestion is performed on the S-Trap column, peptides are eluted and accurately quantified using the fluorescence-based CBQCA method [33]. For total Rab protein analysis, 0.5 μg of tryptic digest (<0.5 % of the total peptides) is spiked with 50 fmol of synthetic heavy labelled isotope Rab1A, Rab1B, Rab3D, Rab8A, Rab8B, Rab10, Rab12, Rab35 and Rab43 markers. Total Rab protein levels are quantified using a 45 min EvoSep LC run and data analyzed as described above. For the immunoprecipitation enrichment step, 150 μg (MEF, neutrophil, mouse lung, kidney, spleen) or 650 μg (mouse brain) derived tryptic peptide digest is spiked with 100 fmol cocktail of pRab1, pRab3, pRab8, pRab10, pRab35, pRab43, LRRK2(388-399), p910 LRRK2, and p935LRRK2 standard heavy labelled peptides. Immunoprecipitations with a mixture of the pan-pRab, 8G10, p910 and p935 antibodies are undertaken. To maximize recovery of pRab43 two rounds of immunoprecipitation are performed. The data is analyzed to quantify the total and phosphorylated levels of pRab1, pRab3, pRab8, pRab10, pRab35 and pRab43 as well as total LRRK2, pSer910-LRRK2 and pSer935-LRRK2.

### Targeted analysis of the LRRK2-Rab pathway in wild type and VPS35[D620N] MEFs

To verify whether our assay can reliably quantify endogenous phosphorylated and total LRRK2 and Rab proteins, we treated littermate-matched, primary wild type and VPS35[D620N] knock-in MEFs with increasing doses of MLi-2 LRRK2 inhibitor and analyzed samples at each dose in triplicate [41]. Cell extracts were prepared and analyzed by either by the targeted MS assay workflow (Fig 4) or by conventional immunoblotting (Fig 5B). Using the targeted MS assay, we were able to quantify all the pRab, total-Rab, total-LRRK2 pSer910/pSer935-LRRK2 endogenous peptides above the linear detection limit (Fig 4). The MS data is presented as fmol peptide/mg of tryptic peptide digest. Total Rab peptides are presented relative to the median intensity of each corresponding Rab peptide (Fig 4). We also analyze the relative fold change between wild type and VPS35[D620N] (Fig 4) and the relative fold change ± 100 nM MLi-2 (Fig 4). MLi-2 IC50 curves for each pRab as well as pSer910/pSer935-LRRK2 are presented in (Fig S6A to H). The data reveal a 5.5-fold increase in pRab10 levels in the VPS35[D620N] compared to wild type MEFs (Fig 4D, Fig 5B). MS analysis demonstrates that MLi-2 reduced pRab10 levels with an IC50 of ~3 nM (Fig 4D, Fig S6D). The VPS35[D620N] mutation increased phosphorylation levels of pRab1 (3.1-fold Fig 4A), pRab3 (2-fold Fig 4B), pRab8 (6.2-fold Fig 4C), pRab35 (3.3-fold Fig 4E) and pRab43 (5.3-fold Fig 4F). The IC_50_ values for inhibition by MLi-2 were 2-3 nM for all of the pRab proteins (Fig: S7A to F). Immunoblotting analysis of the same samples used in the MS analysis confirmed that pRab10 levels were increased around ~6-fold in the VPS35[D620N] knock-in cells and reduced to basal levels in a dose dependent manner by MLi-2 treatment with an IC50 of ~ 3 nM, consistent with previous analysis [15] (Fig 5B).

A high dose of 100 nM MLi-2 reduced pRab10 levels 21-fold in wild type and 75-fold in VPS35[D620N] MEFs (Fig 4D). For wild type cells, pRab43 phosphorylation was reduced over 10-fold by 100 nM MLi-2, with other pRabs impacted between 1.3 to 4-fold (Fig 4). The impact of 100 nM MLi-2 was much greater in VPS35[D620N] knock-in cells displaying higher levels of pRabs. For pRab8, pRab10, pRab35 and pRab43 the effects were over 10-fold, but lower for pRab1 and pRab3 (Fig 4). The relative abundance of the different pRab are shown in Table 1. Ranking abundance of pRabs from high to low in VPS35[D620N] MEFs are pRab10, pRab1, pRab8, pRab35, pRab43 and pRab3. Consistent with previous results the total levels of Rab proteins were similar in wild type and VPS35[D620N] MEFs (Fig 4).

**Table: 1.** Estimated relative abundance of pRabs and LRRK2-pS910, pS935 and total LRRK2 (Fmol/mg)

| Table: 1 | Estimated relative abundance of pRabs and LRRK2-pS910, pS935 and total LRRK2 (Fmol/mg) |  |  |  |  |  |  |  |  |
| --- | --- | --- | --- | --- | --- | --- | --- | --- | --- |
| Sample | pRab1 | pRab3 | pRab8 | pRab10 | pRab35 | pRab43 | pS910<br>LRRK2 | pS935<br>LRRK2 | Total<br>LRRK2 |
| Wt (-MLi2) - MEFs<br>(+MLi2) | 48.1<br>(36.9) | 8.5<br>(5.9) | 34.9<br>(9.2) | 165<br>(7.9) | 20.4<br>(10.5) | 9.9<br>(0.7) | 45.5<br>(4.6) | 53.0<br>(4.5) | 70.6<br>(66.6) |
| LRRK2 R1441C (-MLi2) - MEFs<br>(+MLi2) | 65.7<br>(41.7) | 10.3<br>(4.3) | 127.5<br>(21.1) | 702.7<br>(21.9) | 54.0<br>(13.1) | 31.0<br>(4.3) | 19.6<br>(4.5) | 40.1<br>(8.0) | 47.9<br>(43.6) |
| VPS35 [D620N] (-MLi2) - MEFs<br>(+MLi2) | 171.7<br>(25.0) | 17.1<br>(3.9) | 213.8<br>(6.6) | 798.6<br>(11.8) | 69.3<br>(6.6) | 54.1<br>(3.9) | 40.2<br>(4.5) | 40.1<br>(8.5) | 64.6<br>(57.4) |
| Neutrophils (-MLi2)<br>(+MLi2) | 9.1<br>(8.8) | 16.6<br>(15.3) | 22.7<br>(13.2) | 44.6<br>(6.6) | 16.6<br>(15.3) | 7.1<br>(1.0) | 103.4<br>(5.4) | 88.0<br>(4.1) | 239.7<br>(182.3) |
| NeutrophilsVPS35[D620N] (-MLi2)<br>(+MLi2) | 14.6<br>(11.2) | 23.0<br>(14.9) | 58.6<br>(11.9) | 207.3<br>(13.2) | 20.3<br>(15.1) | 69.8<br>(7.1) | 62.1<br>(14.9) | 148.8<br>(38.0) | 87.9<br>(74.1) |
| Whole brain - Wt (-MLi2)<br>(+MLi2) | 1.5<br>(2.1) | 3.6<br>(3.9) | 0.8<br>(1.6) | 4.2<br>(3.9) | 3.9<br>(1.9) | 0.4<br>(0.3) | 10.3<br>(2.6) | 23.3<br>(1.2) | 24.4<br>(20.8) |
| Whole brain - LRRK2 R1441C (-MLi2) | 3.2<br>(3.0) | 9.0<br>(6.3) | 2.0<br>(1.8) | 14.7<br>(8.1) | 4.1<br>(2.8) | 1.6<br>(0.4) | 3.5<br>(1.5) | 6.3<br>(2.1) | 45.2<br>(42.8) |
| Whole brain - VPS35 [D620N] (MLi2)<br>(+MLi2) | 2.0<br>(2.0) | 4.6<br>(5.5) | 2.1<br>(1.2) | 7.5<br>(4.3) | 3.6<br>(3.8) | 1.2<br>(0.5) | 11.4<br>(4.50) | 17.8<br>(2.0) | 21.8<br>(27.5) |
| Kidney- Wt (-MLi2)<br>(+MLi2) | 3.6<br>(3.8) | 20.4<br>(10.5) | 3.3<br>(1.3) | 13.2<br>(4.0) | 7.9<br>(2.1) | 9.9<br>(1.2) | 69.3<br>(2.3) | 148.5<br>(34.9) | 186.1<br>(215.1) |
| Kideny- LRRK2 R1441C (-MLi2)<br>(+MLi2) | 4.0<br>(3.9) | 22.7<br>(27.8) | 3.3<br>(2.4) | 12.4<br>(5.8) | 19.3<br>(25.7) | 7.7<br>(25.7) | 4.8<br>(2.2) | 9.5<br>(4.0) | 135.5<br>(147.0) |
| Kideny- VPS35 [D620N] (-MLi2)<br>(+MLi2) | 6.0<br>(3.5) | 12.5<br>(11.2) | 5.2<br>(1.9) | 36.3<br>(6.6) | 14.5<br>(5.8) | 19.1<br>(1.1) | 58.0<br>(1.3) | 134.6<br>(27.0) | 215.1<br>(208.0) |
| Lungs- Wt (-MLi2)<br>(+MLi2) | 48.7<br>(47.8) | 21.5<br>(11.7) | 40.0<br>(29.6) | 116.5<br>(30.2) | 17.2<br>(15.2) | 23.9<br>(3.8) | 101.0<br>(5.1) | 331.6<br>(30.8) | 113.1<br>(93.6) |
| Lungs -LRRK2 R1441C (-MLi2)<br>(+MLi2) | 42.2<br>(25.2) | 70.1<br>(63.3) | 53.7<br>(15.0) | 167.1<br>(22.8) | 31.0<br>(5.7) | 37.7<br>(6.4) | 13.0<br>(6.2) | 24.5<br>(8.0) | 94.6<br>(112.7) |
| Lungs [D620N] (-MLi2)<br>(+MLi2) | 106.2<br>(47.0) | 24.5<br>(11.7) | 113.6<br>(44.6) | 352.4<br>(49.1) | 30.4<br>(15.0) | 83.3<br>(4.5) | 114.7<br>(9.5) | 365.0<br>(16.8) | 115.5<br>(98.3) |
| Spleen- Wt (-MLi2)<br>(+MLi2) | 36.0<br>(34.3) | 12.5<br>(19.1) | 28.2<br>(19.8) | 64.1<br>(20.1) | 12.6<br>(9.1) | 123.3<br>(57.3) | 61.9<br>(3.54) | 268.2<br>(13.6) | 130.6<br>(106.6) |
| Spleen- LRRK2 R1441C (-MLi2)<br>(+MLi2) | 24.1<br>(18.4) | 21.3<br>(20.8) | 25.6<br>(13.6) | 79.2<br>(18.1) | 38.8<br>(16.1) | 134.6<br>(45.6) | 8.2<br>(1.3) | 15.3<br>(1.6) | 136.8<br>(135.4) |
| Spleen- VPS35 [D620N] (-MLi2)<br>(+MLi2) | 50.8<br>(35.3) | 15.1<br>(16.8) | 52.1<br>(25.7) | 169.3<br>(30.1) | 17.5<br>(7.8) | 56.0<br>(2.1) | 53.2<br>(9.5) | 247.0<br>(10.4) | 134.1<br>(112.0) |

**Figure 4:**
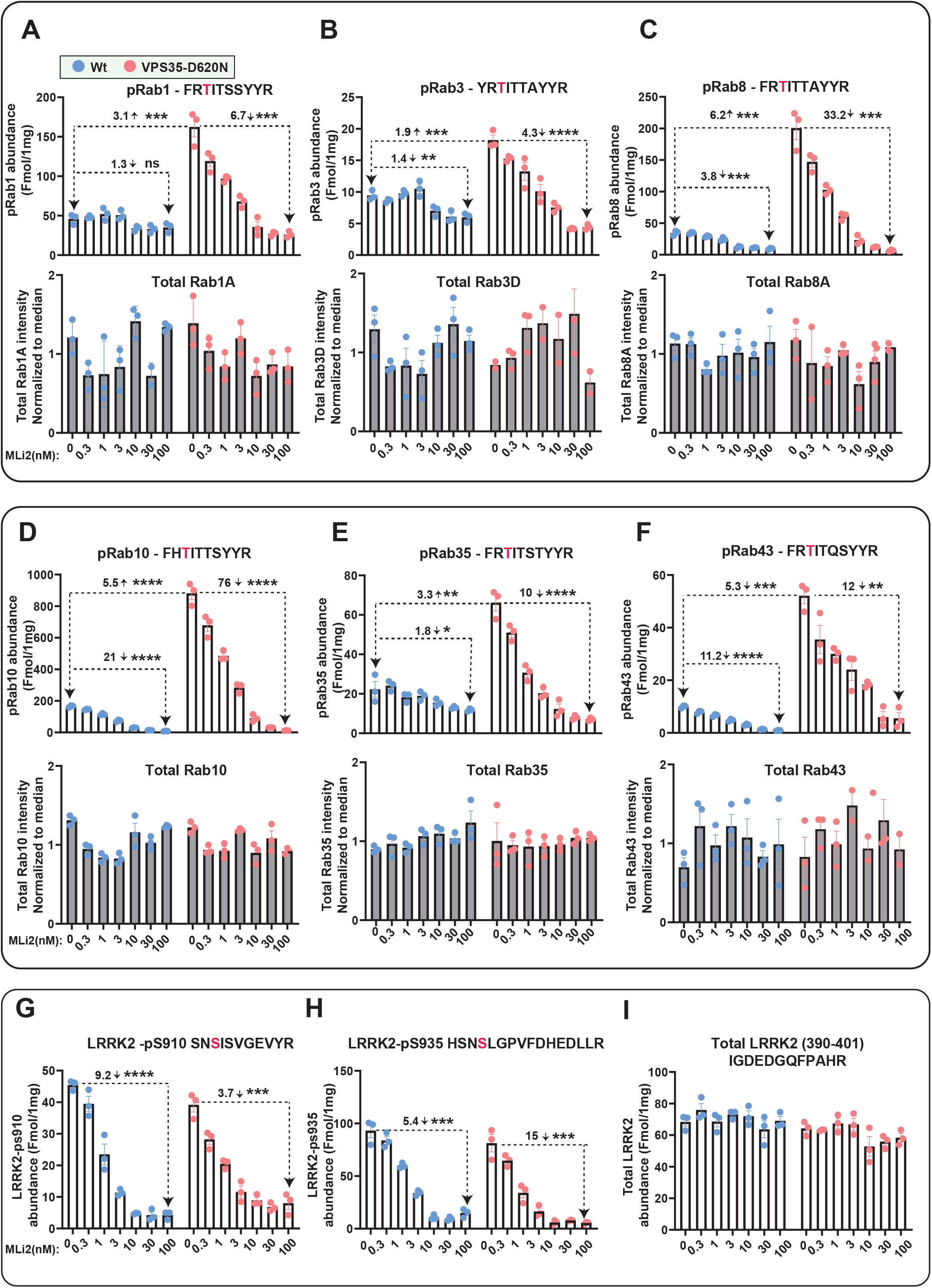
Targeted MS analysis of LRRK2-Rab signalling pathway in wild type and VPS35[D620N] MEFs. Littermate matched wild type and homozygous VPS35 [D620N] MEFs were treated with the indicated concentrations of MLi-2 for 90 min prior to harvest and subjected to either tryptic digestion (**A-I**) or immunoblot analysis (**Figure 5B)**. (**A-I**) 150 μg of digest was spiked with an equimolar ratio of 100 fmol heavy pRabs and LRRK2 pS910/pS935 and total LRRK2 peptides and subjected to sequential immunoprecipitation (IP) with a multiplexed antibody cocktail (Pan-pRab, UDD1, UDD2 and 8G10) and peptide levels quantified as fmol peptide/mg of tryptic digest (upper panel). In parallel, 0.5 μg of tryptic digest spiked with an equimolar ratio of 50 fmol of heavy total Rab peptides and data was acquired in PRM mode and peptide levels normalised to the median peptide intensity (lower panel). Dotted arrows indicate the differences between the indicated samples and one tailed or two tailed student t-Test was performed to depict the significance of change indicated with *. Three independent replicate experiments were performed for each of the indicated MLi-2 concentration and each dot indicates values for each experiment. P values for DMSO versus MLi-2 and WT versus VPS35[D620N] are provided in supplemental table (Table S2). Error bars representing mean ±SEM.

**Figure 5:**
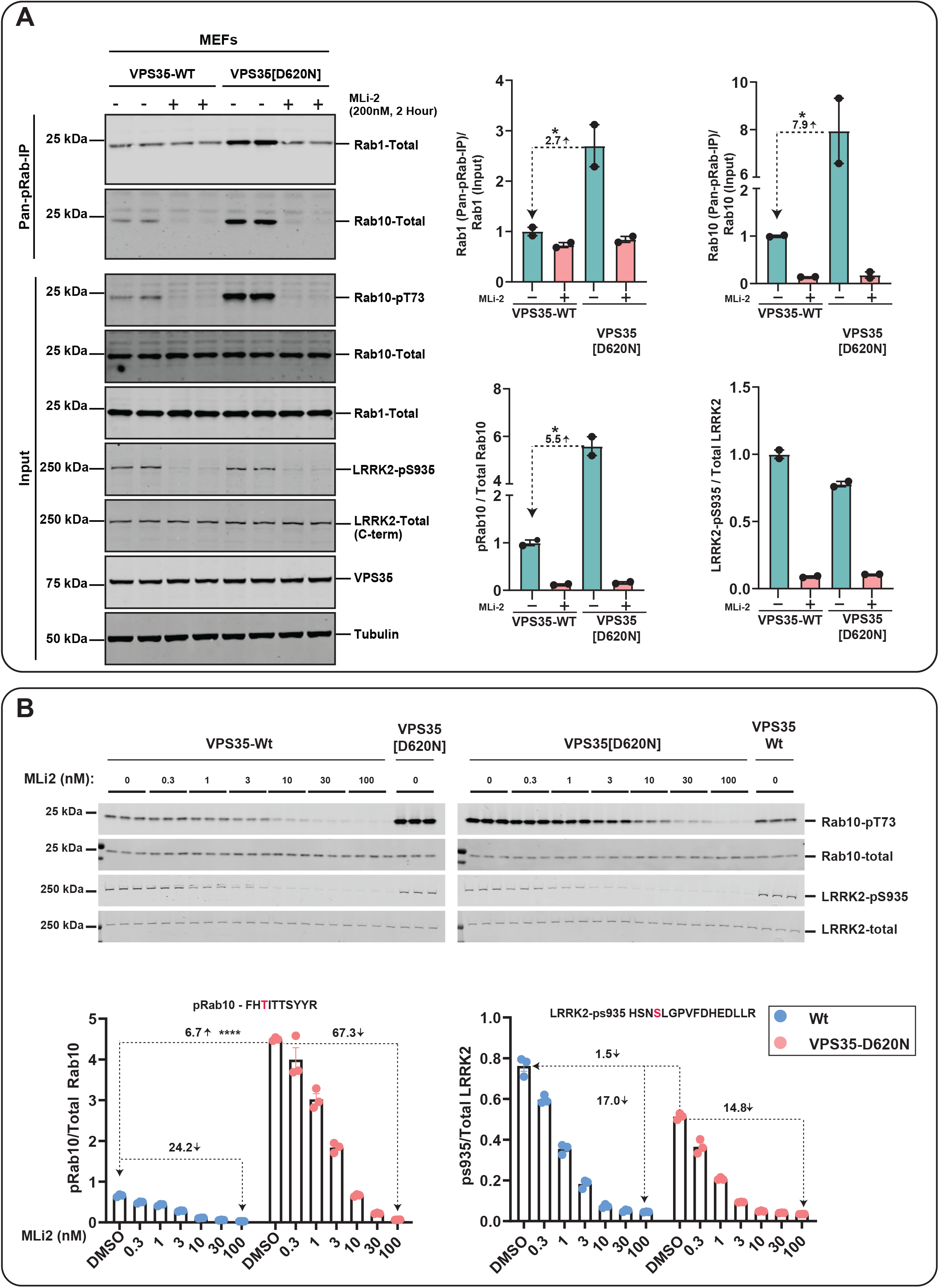
Enhanced LRRK2-mediated Rab1 phosphorylation in homozygous VPS35 [D620N] knock-in MEFs. Littermate matched wild-type and homozygous VPS35[D620N] knock-in mouse embryonic fibroblasts were treated ±200 nM MLi-2 for 2 hours prior to lysis. Immunoprecipitation using the Pan-pRab antibody was performed on 2 mg of whole cell extract and immunoprecipitates were subjected to Rab1-Total (80% of sample) and Rab10-Total (20% of sample) immunoblotting, with 15 μg of whole cell extract used for input (5A). Each lane represents cell extracts obtained from a different dish of cells (two replicates per condition). Immunoblots were subjected to quantitative LI-COR immunoblot analysis with all indicated antibodies at 1 μg/mL, except VPS35 and Tubulin antibodies at 0.1 μg/mL. Bar graphs depicting a significant 2.7 fold elevation of pRab1 in VPS35[D620N] MEFs was observed relative to the wild-type MEFs in Pan-pRab immunoprecipitation similarly a 7.9 fold increase of pRab10 was observed in VPS35[D620N] MEFs. Bar graphs depicting an increase of pRab10 (5.5) fold and LRRK2-pS935 in VPS35[D620N] MEFs input **(B)** The indicated WT and VPS35[D620N] MEFs generated in Figure 4, were treated with or without the indicated concentrations of MLi-2 for 90 min. Cells were lysed, and 10 μg of extract was subjected to quantitative immunoblot analysis with the indicated antibodies (all at 1 μg/ml). Each lane represents cell extract obtained from a different dish of cells (3-4 replicates per condition). The membranes were developed using the Odyssey CLx Western Blot imaging system. Immunoblots were quantified using the Image Studio software. Data are presented relative to the phosphorylation ratio observed in WT cells treated with DMSO (no inhibitor), as mean ± SEM. Statistical significance for pRab10 between VPS35-D620N and WT untreated cells is determined by one tailed T-Test (P=< 0.00001). n=3, error bars representing the mean ±SEM

As outlined in the Introduction, in previous work we had not observed an increase in endogenous pRab1 phosphorylation in LRRK2 pathogenic knock-in MEFs treated ± LRRK2 inhibitors [7]. To obtain further evidence that the VPS35[D620N] mutation increased pRab1 levels, we performed additional immunoblotting of wild type and VPS35[D620N] extracts immunoprecipitated with the pan-pRab antibody. This confirmed that ~3-fold increased levels of Rab1 were immunoprecipitated from VPS35[D620N] compared to wild type cells (Fig 5A). Treatment of VPS35[D620N] cells with ML-2 reduced pRab1 levels to levels observed in wild type cells (Fig 5A). As observed previously [7], and consistent with our MS analysis (Fig 4), MLi-2 did not lower levels of pRab1 in wild type cells (Fig 5A).

MLi-2 also reduced Ser910 and Ser935 phosphorylation in a dose dependent manner, with an IC_50_ of ~ 3 nM in both wild type and VPS35 [D260N] MEFs (Fig 4G,H, Fig 5B). Levels of LRRK2 and pSer910 and pSer935 were identical in wild type and VPS35[D620N] MEFs and not altered by the 90 min MLi-2 treatment (Fig 4G to H), consistent with previous data [15]. These findings are also consistent with immunoblot analysis undertaken in parallel (Fig 5B).

### Validation of targeted MS assay using MLi-2 resistant LRRK2[A2016T] MEFs

We previously described a mutation (A2016T) within the LRRK2 kinase domain of LRRK2 that does not affect basal kinase activity but renders LRRK2 ~10-fold resistant to several kinase inhibitors including MLi-2 [8, 42]. To further validate the targeted-MS assay, we treated littermate-matched, wild type and LRRK2[A2016T] knock-in MEFs [8] with increasing doses of MLi-2 and used targeted MS to calculate abundances of total and phospho Rab proteins as well as total and pSer910/pSer935-LRRK2 (Fig 6); IC_50_ values were also calculated (Fig S8 A-H). For pRab10, pRab8, pRab43, pSer910-LRRK2, pSer935-LRRK2, the A2016T mutation increased IC50 values from 3 nM to 30-40 nM MLi-2 (Fig S8A to C, G to H). MLi-2 had no significant effect on the levels of pRab1, pRab3 or pRab35 in A2016T MEFS (Fig S7A to C). The total levels of Rab and LRRK2 were similar in the wild type and LRRK2 [A2016T] knock-in MEFs (Fig 6A to C and S7A to F). Similar results were obtained in parallel by immunoblotting (Fig S9A to B).

**Figure 6:**
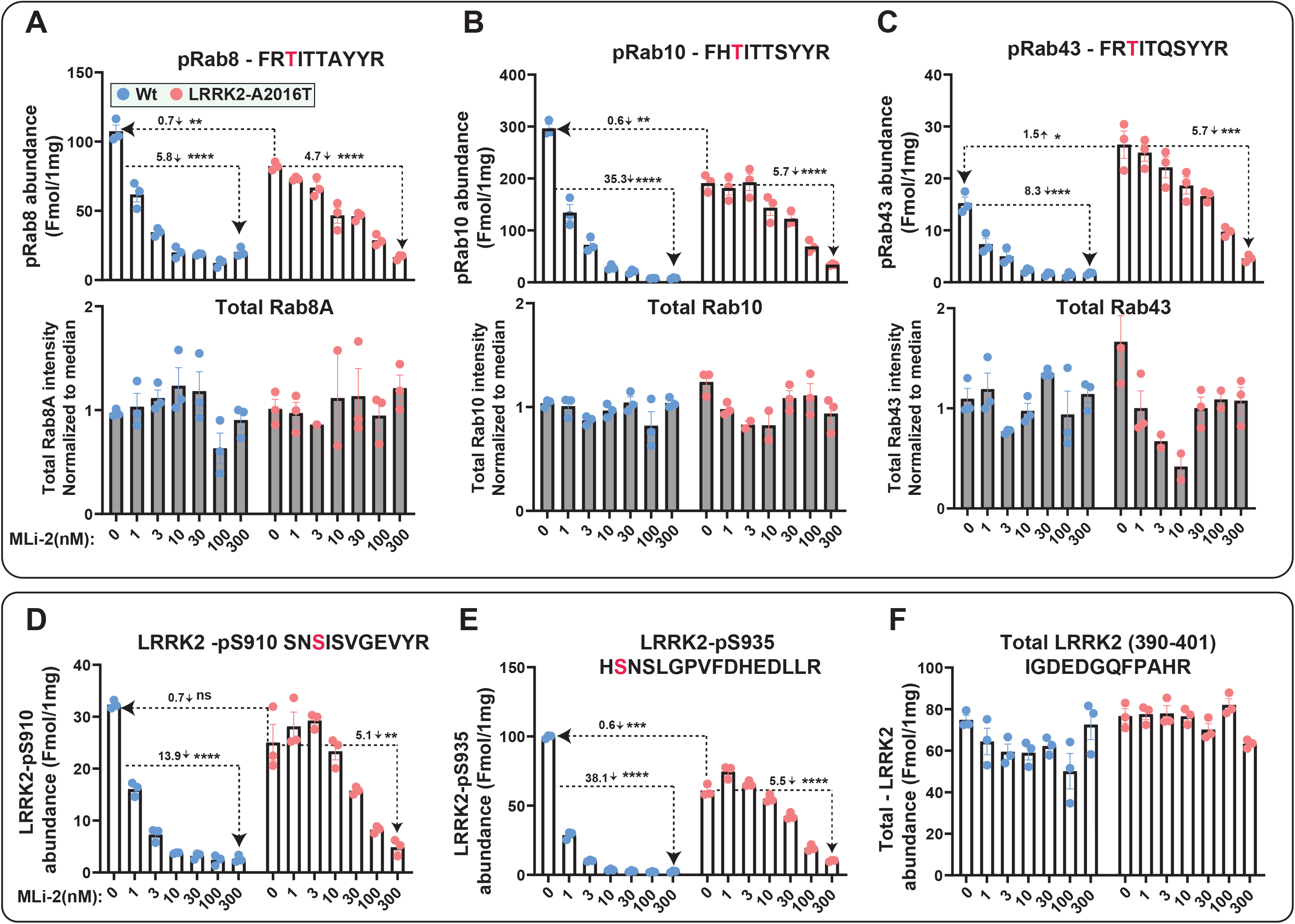
Targeted MS assay of LRRK2-Rab signalling pathway in wild type and LRRK2-A2016T MEFs. Littermate matched wild type and homozygous LRRK2[A2016T] MEFs were treated with the indicated concentrations of MLi-2 for 90 min prior to harvest and subjected to either tryptic digestion (**A-F**) or immunoblot analysis (**Supplementary 9A-B)**. (**A-F**) 150 μg of digest was spiked with an equimolar ratio of 100 fmol heavy pRabs and LRRK2 pS910/pS935 and total LRRK2 peptides and subjected to sequential immunoprecipitation (IP) with a multiplexed antibody cocktail (Pan-pRab, UDD1, UDD2 and 8G10) and peptide levels quantified as fmol peptide/mg of tryptic digest (upper panel). In parallel, 0.5 μg of tryptic digest spiked with an equimolar ratio of 50 fmol of heavy total Rab peptides and data was acquired in PRM mode and peptide levels normalised to the median peptide intensity (lower panel). Dotted arrows indicated the differences between the indicated samples and one tailed or two tailed student t-Test was performed to depict the significance of change indicated with *. Three independent replicate experiments were performed for each of the indicated MLi-2 concentration and each dot indicates values for each experiment. Error bars representing mean ±SEM. Data for pRab1, pRab3, pRab35 that not change significantly with MLi-2 is presented in Figure 7S A-F.

### Further validation of targeted MS assay using LRRK2[R1441C] pathogenic MEFs

We next compared abundance of total and phospho Rab proteins as well as total and pSer910/pSer935-LRRK2 levels in littermate-matched wild type and LRRK2[R1441C] pathogenic mutant MEFs treated ± 100 nM MLi-2 using the targeted MS assay (Fig 7). This revealed significantly increased levels of pRab8 (2.7-fold), pRab10 (2.5-fold), pRab35 (2.6-fold) and pRab43 (3.5-fold) in the LRRK2[R1441C] knock-in MEFs compared with wild type. MLi-2 treatment markedly reduced pRab and pSer910/pSer935-LRRK2 levels (Fig 7A). The LRRK2 [R1441C] mutation had little impact on pRab1 or pRab3 levels (Fig 7B). Previous studies have shown that the LRRK2[R1441C] mutation reduces phosphorylation of LRRK2 at Ser910 and Ser935 [22]. Consistent with this, the targeted MS assay revealed a 2-fold reduction in the phosphorylation of Ser910 and Ser935 phosphorylation sites in the R1441C MEFs compared to wild type (Fig 7A,B). Similar results were also observed in parallel immunoblotting results for pRab10 and pSer935-LRRK2 (Fig7C,D).

**Figure 7:**
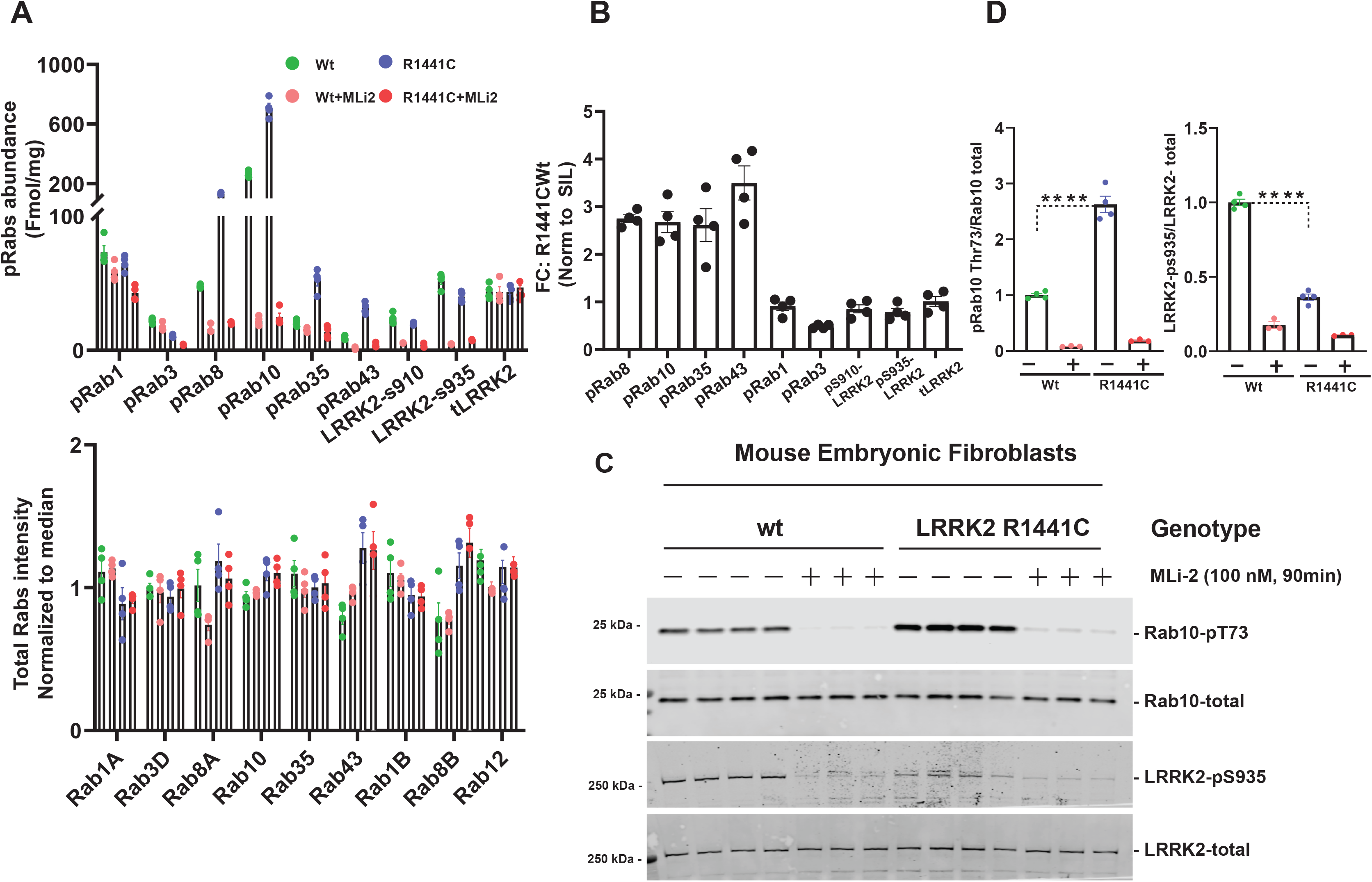
Targeted MS assay of LRRK2-Rab signalling pathway in wild type and LRRK2-R1441C MEFs. Littermate matched wild type and homozygous LRRK2[R1441C] MEFs were treated ± 100 nM MLi-2 for 90 min prior to harvest and subjected to either tryptic digestion (**A,B**) or immunoblot analysis (**C**). (**A**) 150 μg of digest was spiked with an equimolar ratio of 100 fmol heavy pRabs and LRRK2 pS910/pS935 and total LRRK2 peptides and subjected to sequential immunoprecipitation (IP) with a multiplexed antibody cocktail (Pan-pRab, UDD1, UDD2 and 8G10) and peptide levels quantified as fmol peptide/mg of tryptic digest (upper panel). In parallel, 0.5 μg of tryptic digest spiked with an equimolar ratio of 50 fmol of heavy total Rab peptides and data was acquired in PRM mode and peptide levels normalised to the median peptide intensity (lower panel). Dotted arrows indicated the differences between the indicated WT and R1441c samples and two tailed student t-Test was performed to depict the significance of change indicated with *. Four independent replicate experiments were performed for each of the indicated MLi-2 concentration and each dot indicates values for each experiment. P values for WT versus LRRK2[R1441C] are provided in supplemental table 2. Error bars representing mean ±SEM. (**B**) Bar chart indicating the relative fold change between levels of indicated peptides between LRRK2[R1441C] versus wild type ± SEM. (**C**) 10 μg of extract was subjected to quantitative immunoblot analysis with the indicated antibodies (all at 1 μg/ml). Each lane represents cell extract obtained from a different dish of cells (3-4 replicates per condition). The membranes were developed using the Odyssey CLx Western Blot imaging system. Immunoblots were quantified using the Image Studio software. Data are presented relative to the phosphorylation ratio observed in WT cells treated with DMSO (no inhibitor), as mean ± SEM. One tailed student t-Test representing the statistical significance of pRab10 and LRRK2-pS935 between LRRK2 R1441C and WT MEFs. Significance representing for the immunoblot analysis of pRab10 (p= 0.000036), LRRK2-pS935 (<0.00001). Error bars indicating the mean ± SEM.

### LRRK2-Rab targeted MS assay on wild type and LRRK2[R1441C] mouse tissues

We next evaluated whether the targeted MS assay could be deployed to assess the LRRK2-Rab pathway in tissues extracts derived from littermate wild type and LRRK2[R1441C] Knock-in mice (Fig 8). For these studies we analyzed 6 mice per group, two of which were treated by subcutaneous injection of 30 mg/kg MLi-2 for 2 hours prior to tissue harvesting. We undertook tissue analysis from kidney and whole brain, lung and spleen (Fig 8) in which the LRRK2 signalling pathway has been investigated previously [21]. For lung, kidney and spleen, 150 μg of tryptic peptide digest was used and was sufficient to detect robustly all pRabs and total-LRRK2 and pSer910/pSer935-LRRK2. For the brain, pRab levels were around 4-fold lower, and we therefore utilized 600 μg of tryptic peptide digest derived from 1 mg cell extract. The relative abundance of each pRab protein in every tissue ± MLi-2 is shown in Figure 8 and Table 1. The LRRK2[R1441C] mutation moderately increased Rab10 phosphorylation in brain (1.5-fold), lung (1.7-fold), kidney (1.7-fold) and spleen (1.5-fold). In the brain, the LRRK2[R1441C] mutation also increased Rab43 phosphorylation 3-fold. In lung, kidney and spleen pRab10, pRab35 and pRab43 were the pRabs most increased by the R1441C mutation (Table 1). MLi-2 administration significantly reduced these pRabs to lower levels similar to levels observed in wild type mice treated with MLi-2 (Fig 8, Table 1). Consistent with previous work [22], levels of Ser910 and Ser935 phosphorylated LRRK2 were reduced 4 to 5-fold in all tissues from LRRK2[R1441C] animals compared to wild type. Similar results were also observed in parallel immunoblotting results for pRab10 and pSer935-LRRK2 (Fig S10A to B).

**Figure 8:**
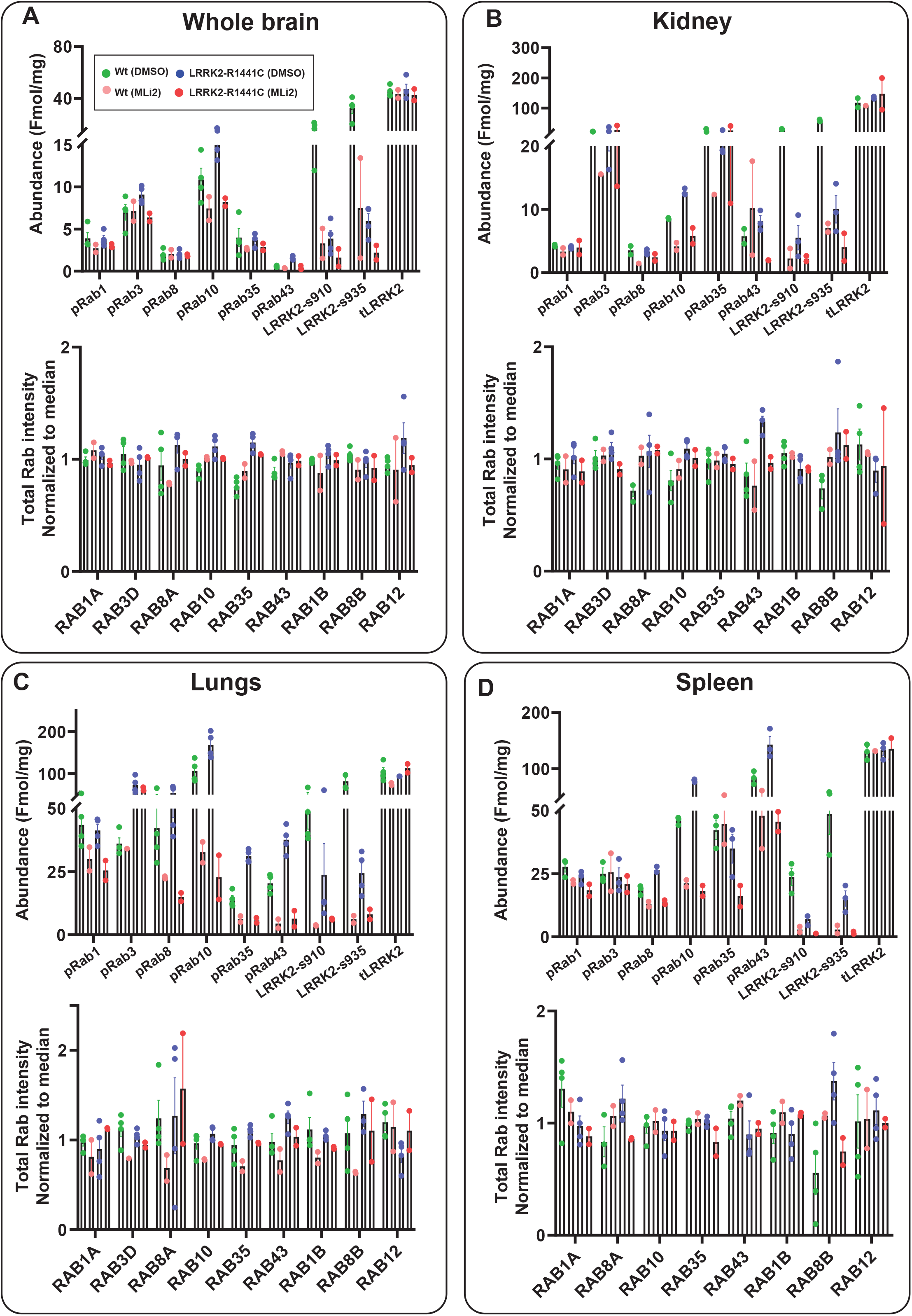
Targeted MS assay of LRRK2-Rab signalling pathway in WT and LRRK2[R1441C] mouse tissues: (A-D) Wild type and homozygous LRRK2 R1441C litter mate matched six months old mice were administered with vehicle (40% (w/v) (2-hydroxypropyl)-β-cyclodextrin) or 30 mg/kg MLi-2 dissolved in vehicle was injected sub-cutaneous 2 hours prior to the tissue collection. Tissues were harvested and subjected to either tryptic digestion (**A to D**) or immunoblot analysis (**Fig S10 A-B**). (**A to D**) Either 600 μg (brain) or 150 μg (lung, kidney, spleen) tryptic digest was spiked with an equimolar ratio of 100 fmol heavy pRabs and LRRK2 pS910/pS935 and total LRRK2 peptides and subjected to sequential immunoprecipitation (IP) with a multiplexed antibody cocktail (Pan-pRab, UDD1, UDD2 and 8G10) and peptide levels quantified as fmol peptide/mg of tryptic digest (upper panel). In parallel, 0.5 μg of tryptic digest spiked with an equimolar ratio of 50 fmol of heavy total Rab peptides and data was acquired in PRM mode and peptide levels normalised to the median peptide intensity (lower panel). 2 to 4 independent replicate experiments were performed for each of the indicated conditions and each dot indicates values for each experiment. Error bars representing mean ±SEM.

### Use of the LRRK2-Rab targeted MS assay on wild type and VPS35[D620N] knockin mouse tissues

We further analyzed LRRK pathway components in brain, lung, kidney and spleen from littermate-matched wild type and VPS35[D620N] mice, also using 6 animals in each group (4 untreated and 2 treated with 30 mg/kg MLi-2 for 2 h). VPS35[D620N] mutation enhanced pRab protein phosphorylation to a greater extent than observed with the R1441C mutation (compare Figs 7, 8). In brain, the VPS35[D620N] enhanced pRab10 (1.8-fold), pRab8 (1.6-fold), Rab43 (3-fold) with no change observed for pRab1, pRab3 and pRab35 (1.4-fold). In other tissues, the VPS35[D620N] mutation increased pRab10, pRab8, Rab43 and pRab35 1.3 - 3.1-fold (Fig 9). As in MEFs, the D620N mutation enhanced pRab1 phosphorylation 1.3 - 1.9-fold in lung, kidney and spleen. MLi-2 reduced levels of pRab8, pRab10, pRab35 and pRab43 as well as pRab1 in VPS35[D620N] MEFs, 2 to 10-fold without impacting pRab3. Levels of Ser910 and Ser935 phosphorylated LRRK2 were similar in wild type and VPS35[D620N] tissues. Analogous results were also observed in parallel immunoblotting results for pRab10 and pSer935-LRRK2 (Fig S11A to B).

**Figure 9:**
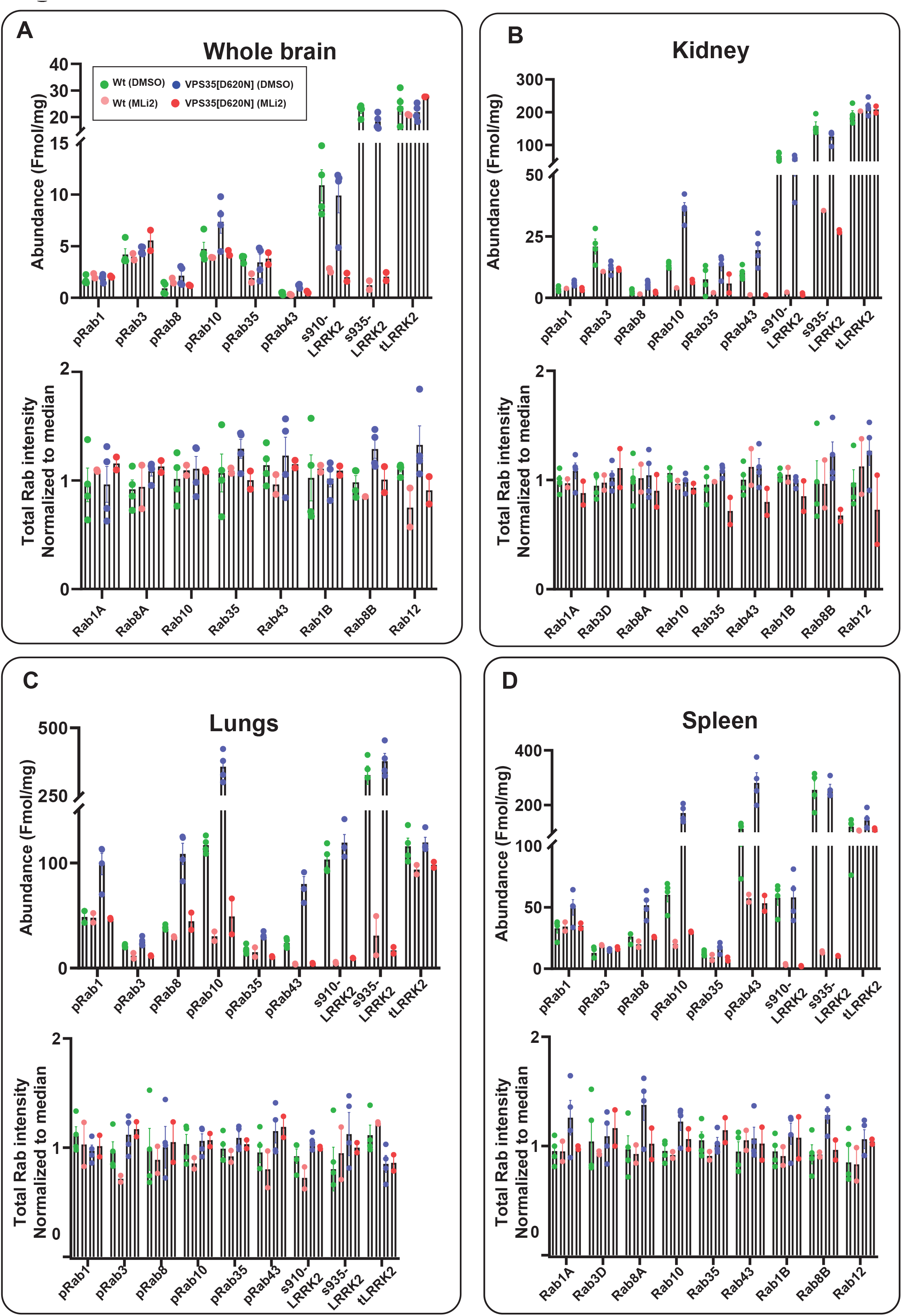
Targeted MS assay of LRRK2-Rab signalling pathway in WT and VPD35[D620N] mouse tissues: (A-D) Wild type and homozygous VPS35[D620N] litter mate matched six months old mice were administered with vehicle (40% (w/v) (2-hydroxypropyl)-β-cyclodextrin) or 30 mg/kg MLi-2 dissolved in vehicle was injected sub-cutaneous 2 hours prior to the tissue collection. Tissues were harvested and subjected to either tryptic digestion (**A to D**) or immunoblot analysis (Fig 11S A-B). (**A to D**) Either 650 μg (brain) or 150 μg (lung, kidney, spleen) tryptic digest was spiked with an equimolar ratio of 100 fmol heavy pRabs and LRRK2 pS910/pS935 and total LRRK2 peptides and subjected to sequential immunoprecipitation (IP) with a multiplexed antibody cocktail (Pan-pRab, UDD1, UDD2 and 8G10) and peptide levels quantified as fmol peptide/mg of tryptic digest (upper panel). In parallel, 0.5 μg of tryptic digest spiked with an equimolar ratio of 50 fmol of heavy total Rab peptides and data was acquired in PRM mode and peptide levels normalised to the median peptide intensity (lower panel). 2 to 4 independent replicate experiments were performed for each of the indicated conditions and each dot indicates values for each experiment. Error bars representing mean ±SEM.

### Use of the LRRK2-Rab PRM MS assay on human primary neutrophils

In clinical studies, the LRRK2 pathway has been evaluated in primary neutrophils as these cells express high levels of LRRK2 and can be easily isolated from donors and patients [15, 30, 31]. To first evaluate the utility of the targeted MS assay, neutrophils were isolated from 3 healthy donors and cells treated with increasing concentrations of MLi-2 prior to lysis. Similar to previous observations [25, 30], immunoblotting analysis revealed that MLi-2 markedly reduced pRab10 levels at 100-300 nM with little inhibition observed at lower concentrations (Fig S12). Similar results were observed for pRab10 andpSer910/pSer935-LRRK2 (Fig S12). MLi-2 (100 or 300 nM) reduced pRab10, pRab43, pSer910 and pSer935 by over 80%. In contrast, pRab1, pRab3, pRab8 and pRab35 were not reduced by MLi-2 administration. We observed some variation in pRab levels between donors. For example, donor-2 displayed moderately higher total LRRK2, pRab10 and pSer910/pSer935-LRRK2 (Fig S12). Similar results were also observed in parallel immunoblotting results for pRab10 and pSer935-LRRK2 (Fig S12E, F).

Finally, we obtained neutrophils form 3 Parkinson’s patients with VPS35[D620N] mutations and 3 healthy control donors (Fig 10). Previous work by immunoblot [15] and MS analysis [25], indicated that neutrophils isolated from VPS35[D620N] subjects displayed 2 to 4-fold elevation in pRab10 levels. Consistent with previous work, our targeted MS assay reveals 2 to 5.4-fold elevation in pRab10 levels in the 3 sets of neutrophils analyzed compared to controls (Fig 10A). Interestingly, pRab43 (Fig 10B) and pRab8 (Fig 10C) were also elevated up to 6-fold in VPS35[D620N] subjects. We also noted a moderate ~1.5 fold elevation of pRab1 (Fig 10D) and pRab3 (Fig 10E) in the VPS35[D620N] subjects. In contrast, levels of pRab35 were barely increased in the VPS35[D620N] neutrophils (Fig 10F). Immunoblot analysis (Fig 10J) of these samples reveals strong correlation with MS analysis between the subjects. The subjects display the equivalent degree of elevated pRab10 levels in both assays (compare Fig 10A with 10J).

**Figure 10:**
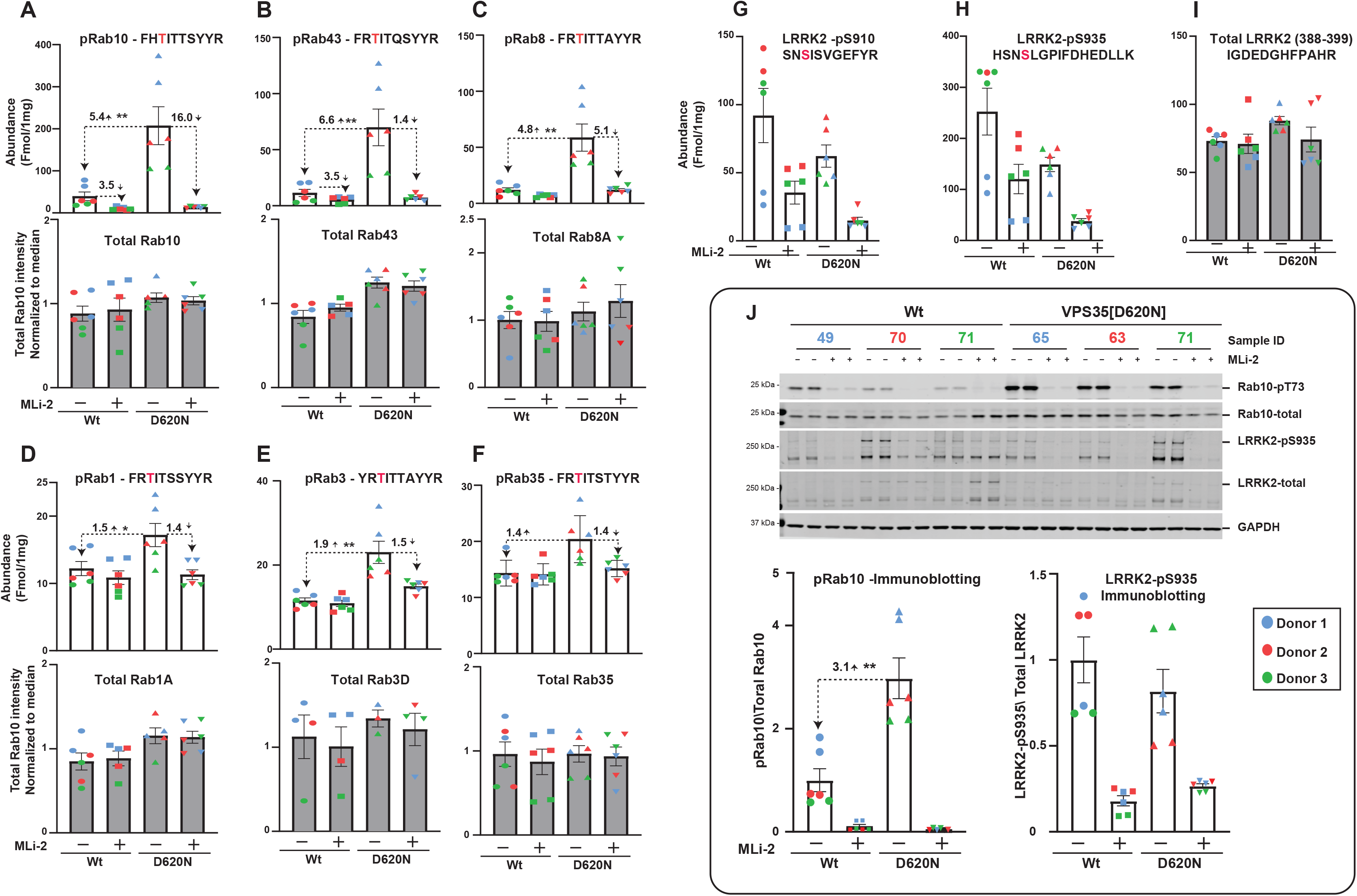
Targeted MS analysis of LRRK2-Rab signalling pathway in healthy donors and VPS35 [D620N] patients neutrophils: (**A-I**) 60 μg of tryptic digest as technical duplicates from 3 healthy donors and 3 VPS35 [D620N] patient samples that are treated with ±MLi-2 was spiked with an equimolar ratio of 100 fmol heavy pRabs and LRRK2 pS910/pS935 and total LRRK2 peptides and subjected to sequential immunoprecipitation (IP) with a multiplexed antibody cocktail (Pan-pRab, UDD1, UDD2 and 8G10) and peptide levels quantified as fmol peptide/mg of tryptic digest (upper panel). In parallel, 0.5 μg of tryptic digest spiked with an equimolar ratio of 50 fmol of heavy total Rab peptides and data was acquired in PRM mode and peptide levels normalised to the median peptide intensity (lower panel). Dotted arrows indicate the differences between the indicated samples and one tailed or two tailed student t-Test was performed to depict the significance of change indicated with *. Three independent replicate experiments were performed for each of the indicated MLi-2 concentration and each dot indicates values for each experiment. Error bars representing mean ±SEM. **(J)** 10 μg of extract was subjected to quantitative immunoblot analysis with the indicated antibodies (all at 1 μg/ml). Each lane represents cell extract obtained from a different donors colour coded (Donor1=blue, Donor2=red and Donor3= green). Data are presented relative to the phosphorylation ratio observed in cells treated with DMSO (no inhibitor), as mean ± SEM. Statistical analysis was performed by two tailed student T-test indicating the significance between VPS35 [D20N] and healthy donors.

## Discussion

The LRRK2-Rab targeted MS assay described here requires a hybrid quadrupole Orbitrap mass spectrometry instrument and preferably an EvoSep liquid chromatography system. The key antibodies (Pan-Rab, 8G10, UDD1 and UDD2) used for the peptide enrichment immunoprecipitation step are commercially available and the series of heavy labeled, stable synthetic peptides (S. Table 1) required for accurate identification and quantification can be easily obtained. For cells and tissues that express relatively high levels of LRRK2 (MEFs, mouse lung, kidney, spleen and human neutrophils), 150 μg of tryptic peptide digest that can readily be obtained from 0.2-0.25 mg of cell/tissue extract is sufficient for analysis. For neutrophil samples derived from Parkinson’s patients for which sample volume was limiting we used 100 μg of cell extract and 60 μg of tryptic peptide digest. For brain samples where levels of LRRK2 are expressed at lower levels, we utilized 600-650 μg of tryptic peptide digest that can be obtained from 0.8 to 1 mg of brain extract. In all cells and tissue extracts analyzed, we detected and quantified six LRRK2-phosphorylated pRab proteins (pRab1, pRab3, pRab8, pRab10, pRab35, pRab43) as well as the total Rab, total LRRK2 and pSer910/pSer935 peptides. The levels of these components are comfortably above the assay linear limit of detection. We present the abundance levels of these peptides as fmol detected peptide/μg of tryptic peptide digest that permits the relative levels of pRab proteins to be compared between different tissues and cells (Table 1).

The S-Trap on column digestion workflow allows for a simplified tryptic peptide digest preparation without requiring an additional desalting step. The tryptic digestion and immunoprecipitation steps could be automated using liquid handling platforms to increase throughput of the assay in a 96 well plate format. Sample preparation to the MS requires 2 days (1 day for peptide digest and 1 day for immunoprecipitation and sample clean up). To prepare 24 samples for tryptic peptide digest requires ~8 h on day 1 (2h reduction and alkylation, 2h S-Trap loading and clean up, 2 hours tryptic digestion, 2h peptide quantitation) followed over overnight lyophilization. Day 2 requires 9h (2x 2h antibody incubation, 2h washing immunoprecipitates, 2h desalting samples and 1 h vacuum drying). Samples at this stage can stored at −80 °C at this stage or directly loaded on the Evotips (2h for 24 samples) and then stored at 4 °C for up to 1 week prior to MS analysis. For the total Rab samples, aliquots of the tryptic peptides are directly loaded on the Evotips. The MS instrument time required to analyze the pRabs, pSer910/pSer935 and total LRRK2 is 21 min, enabling ~60 samples to be processed per day. The total Rab samples require a longer 45 min gradient, allowing ~ 30 samples per day to be analyzed. If both pRabs and total Rab peptides are evaluated in parallel, ~20 sets of samples can be analyzed per day. We find that the levels of total Rabs are constant between cell lines and tissues, and if dose dependence curves and time courses are being undertaken, it may not be necessary to monitor total levels of Rab protein for every dose or time point.

One of the key limitations of our targeted MS assay is that it does not detect pRab12, as this peptide is not immunoprecipitated with the Pan-pRab antibody (FIG XX). pRab12 levels are especially high in brain and are more impacted than Rab10 by LRRK2 mutations and inhibitors in that tissue [21, 43]. Only a single pRab12 phospho-specific monoclonal antibody is commercially available and we have found that this does not immunoprecipitate pRab12 peptide (Figure S5D, E). In future work it would be important to raise an immunoprecipitating pRab12 antibody to enable pRab12 levels to be quantified. Our studies are consistent with previous work [21, 43], that in wild type brain, pRab levels including Rab10 are lower compared to other tissues and cells analyzed and MLi-2 administration has little impact on any of the impact on the phosphorylation levels of any other Rab protein studied (Fig 8A, 9A).

In Table 1 we summarize our data on the relative levels of pRab proteins in MEFs, human neutrophils, mouse brain, lung, kidney and spleen. In MEFs, neutrophils, and lung, pRab10 is abundant. In the kidney and spleen, levels of pRab3 and pRab43, respectively, are moderately higher than pRab10. The fold-impact of MLI-2 is greatest for the pRabs whose phosphorylation is most enhanced by the LRRK2[R1441C] or VPS35[D620N] mutation. We observed that the VPS35[D620N] mutation in MEFs, as well as lung and kidney, increased pRab1 phosphorylation 1.6 to 3.5-fold in a manner that was suppressed by MLi-2. In MEFs we confirmed this result by immunoblotting analysis (Fig 5A). Consistent with previous work [7], pRab1 levels are relatively high in wild type cells and tissues, but not elevated further in LRRK2[R1441C] pathogenic samples or lowered following treatment with MLi-2 (Table 1). Rab1 is reportedly phosphorylated by TAK1 kinase at this LRRK2 site (Thr75) [44]. Future work is required to uncover the mechanism by which the VPS35[D620N] mutation promotes LRRK2-mediated Rab1 phosphorylation as well as other its other Rab substrates. It is possible that the VPS35[D620N] mutation may enhance the localization of LRRK2 at a membrane compartment containing Rab1. Our data with a small number of neutrophils isolated from VPS35[D620N] Parkinson’s patients demonstrates a modest ~1.5-fold increase in pRab1 levels that are reduced with MLI-2 (Fig 10D). In addition to marked elevation of pRab10 phosphorylation in VPS35[D620N] neutrophils, we observed up to 6-fold increases in pRab8 (Fig 10C) and pRab43 (Fig 10B). It would be important to extend this work with a greater number of subjects in future studies.

In conclusion, we describe a versatile new assay that enables the relative quantification of the key components of the LRRK2 pathway, namely pRabs, pSer910/pSer935-LRRK2 as well as total Rab proteins and LRRK2 to be robustly quantified in cells and tissues. We demonstrate that this assay also works well to monitor the LRRK2 pathway in human neutrophils and shows strong suitability to analyze clinical samples. In our view, this assay provides significantly richer information than can be obtained by immunoblotting in terms of how modulation of LRRK2 signalling pathway influences the phosphorylation of different Rab proteins.

## Acknowledgements

We thank Paul Davies for helpful discussion and epitope mapping the LRRK2 8G10 antibody, Alexia Kalogeropulou for supporting the mouse tissue studies, the excellent technical support of the MRC-Protein Phosphorylation and Ubiquitylation Unit (PPU) mass spectrometry team (coordinated by Renata Soares), Cloning team (coordinated by Rachel Toth), DNA Sequencing Service (coordinated by Gary Hunter), the MRC-PPU tissue culture team (coordinated by Edwin Allen), MRC PPU Reagents & Services antibody and protein purification teams (coordinated by Hilary McLauchlan and James Hastie).

## Author Contributions

R.S.N designed, executed all experiments in this study presented in Figures 1–10, S1–S12, analyzed and interpreted data and helped write the manuscript. F.T. designed and executed experiments shown in Figures 5 B, 7C, 10J, S9 A,B; S10A, B, and S11-A,B and S12-E. And analyzed data. M.T. performed experiments shown in Figure 5A and S5 C. P.L performed experiments shown in Figure 5A and S2 A, S5 C. A.Z and E.S helped with procurement of neutrophils samples. D.R.A. helped with experimental design, analysis and interpretation of data and preparation of the manuscript.

---

**Supplementary Figure 1:**
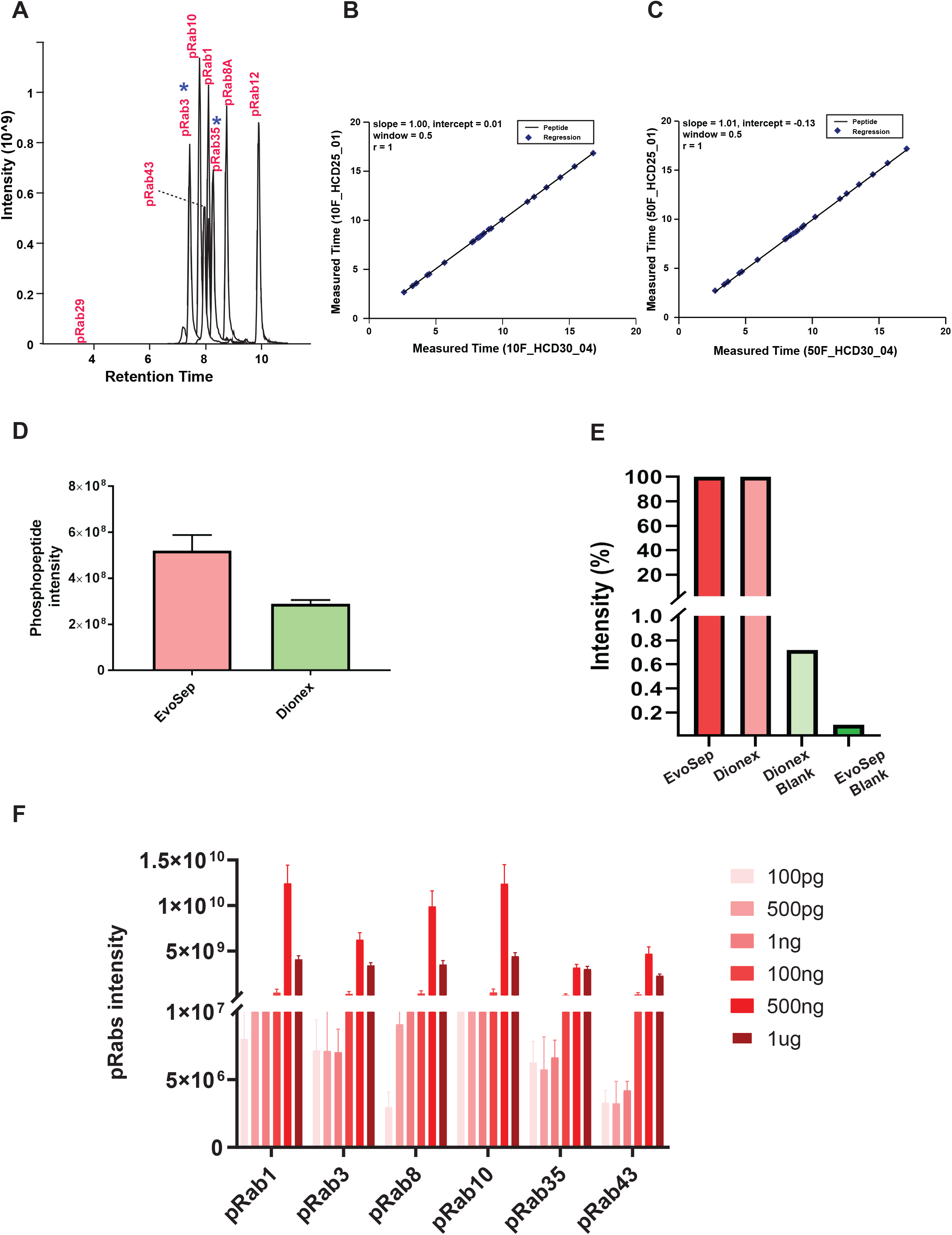
S-**1A)** Equimolar ratio of the indicated pRab peptides are spiked into 50 ng of HeLa digest and analysed using EvoSep 21 min run gradient on QE HF-X MS. Extracted ion chromatograms depicting the separation of pRab peptides on a 21 min run gradient. pRabs are well separated between 7 to 8.5 retention time including the isobaric pRab3 and pRab35 that are well separated by 1 min apart indicated with blue coloured asterisk (*).**S1B-C)**) The retention times reproducibility was superior on 21 min run EvoSep LC system. 10fmol(S1-B) or 50fmol (1S-1C) pRabs measured retention times depicted as a linear regression curve showing an r value closer to 1. **S1-D**) 50 fmol of pRab10 synthetic peptide was analysed using 21 min on EvoSep LC system and 45 min on Dionex RSLC nano LC system. Data was acquired in a PRM mode on QE HF-X instrument. The data was analysed using Skyline software and the relative pRab10 intensity was depicted as a bar graph. Y-axis indicating the pRab10 intensity and x-axis showing the data acquired on EvoSep and Dionex LC systems. **S1-E**) the data acquired in S1-D was assessed for the carryover between the two LC-systems by measuring the pRab10 intensities in blank runs. Carryover appeared to below 0.01% on EvoSep LC system. **S1-F)** Sensitivity of the Pan-pRab antibody was assessed at a varied amounts ranging between 0.001 μg to 1 μg. 100 fmol of pRabs heavy labelled peptide mixture was spiked in to 200 μg of BSA digest and Immunoprecipitation was undertaken with an indicated antibody amounts. Bar graph depicting the relative pRabs intensities on y-axis for each of the pRab peptide on x-axis. The varied amount of antibody was highlighted in a light to dark red coloured gradient. n=3, Error bars representing the mean ±SEM.

**Supplementary Figure 2:**
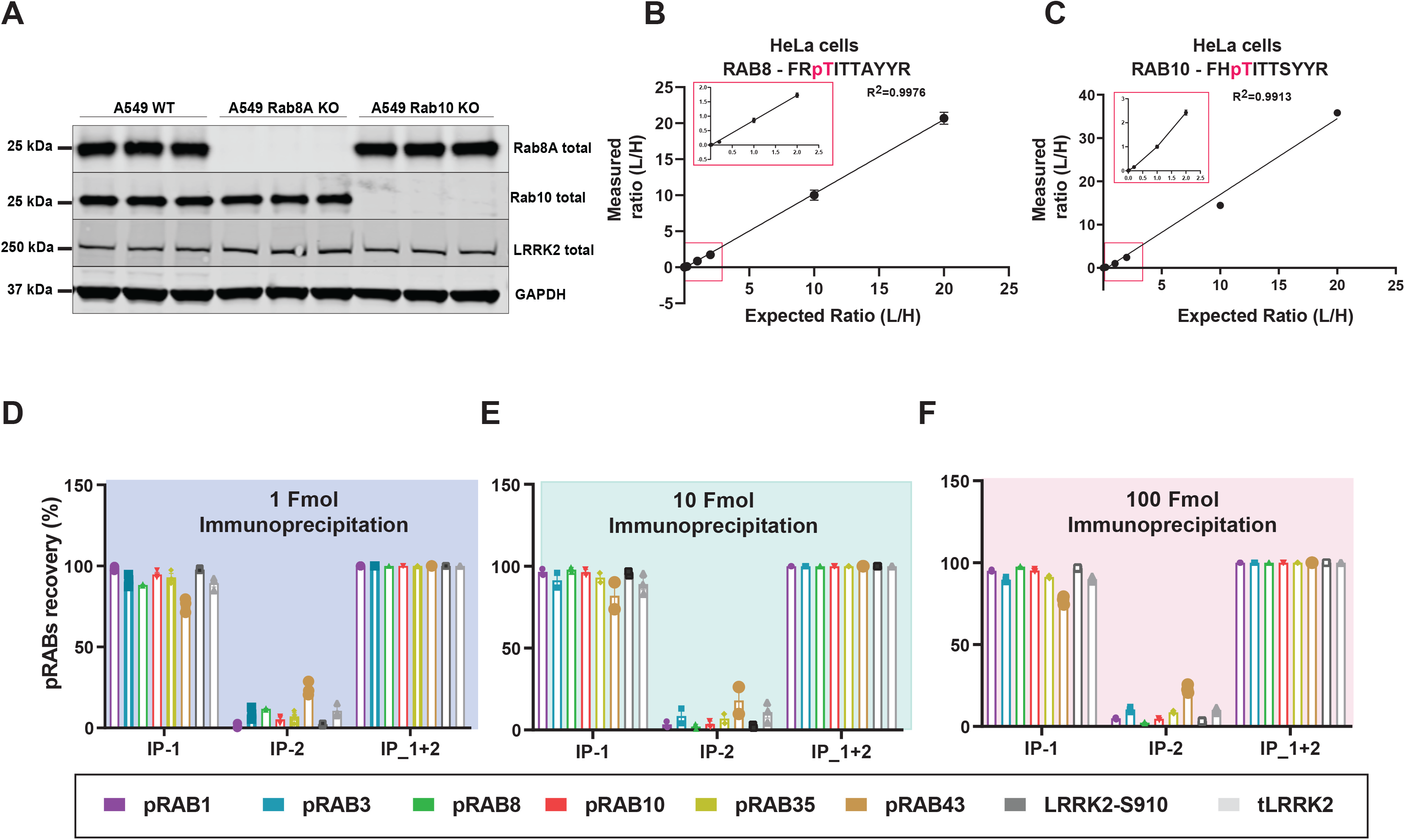
Limit of detection and quantification of targeted pRab8 and pRab10 peptides: **S2-A)** Immunoblotting Validation of Rab8-KO and Rab10-KO A549 cells used in Figure 2A–2B to assess the limit of detection. **S2B-S2C)** Limit of detection experiments were carried out by spiking synthetic light phosphorylated Rab8A (Thr72) peptide at (0.01, 0.1 1, 10, 50, 100, 500, 1000 fmol), while keeping the constant 50 fmol of heavy phosphorylated Rab8A (Thr72) peptide in 50ng of HeLa peptide digest. Immunoprecipitations were undertaken with Pan-pRab antibody and analysed on Orbitrap HF-X MS in a targeted PRM mode. XIC were generated using Skyline software. X-axis depicting the expected (L/H) ratio and Y-axis depicting measured (L/H) ratio. Rectangular box indicates the zoom in of 0.01, 0.1 and 1 fmol data points. n=3. **S2-B)** similar experiments were carried out for Human specific LRRK2 peptides. Error bars represent the mean ±SEM. **S2: D-F)** as described in Figure 2C–2E, similar experiments were carried out for Human specific LRRK2 peptides. The relative pRabs and pLRRK2 and total LRRK2 peptides in ach round of IP was plotted as bar graphs. X-axis representing the pRabs or LRRK2 and Y-axis representing the peptide recovery percentage. Similar experiments were undertaken by spiking 1 fmol (S2-D), 10 fmol (S2-E) and 100 fmol (S2-F) amounts. pRab43 recovery was found to be ~70% and ~30% in IP1 and IP2. n=3, error bars representing mean ±SEM

**Supplementary Figure 3:**
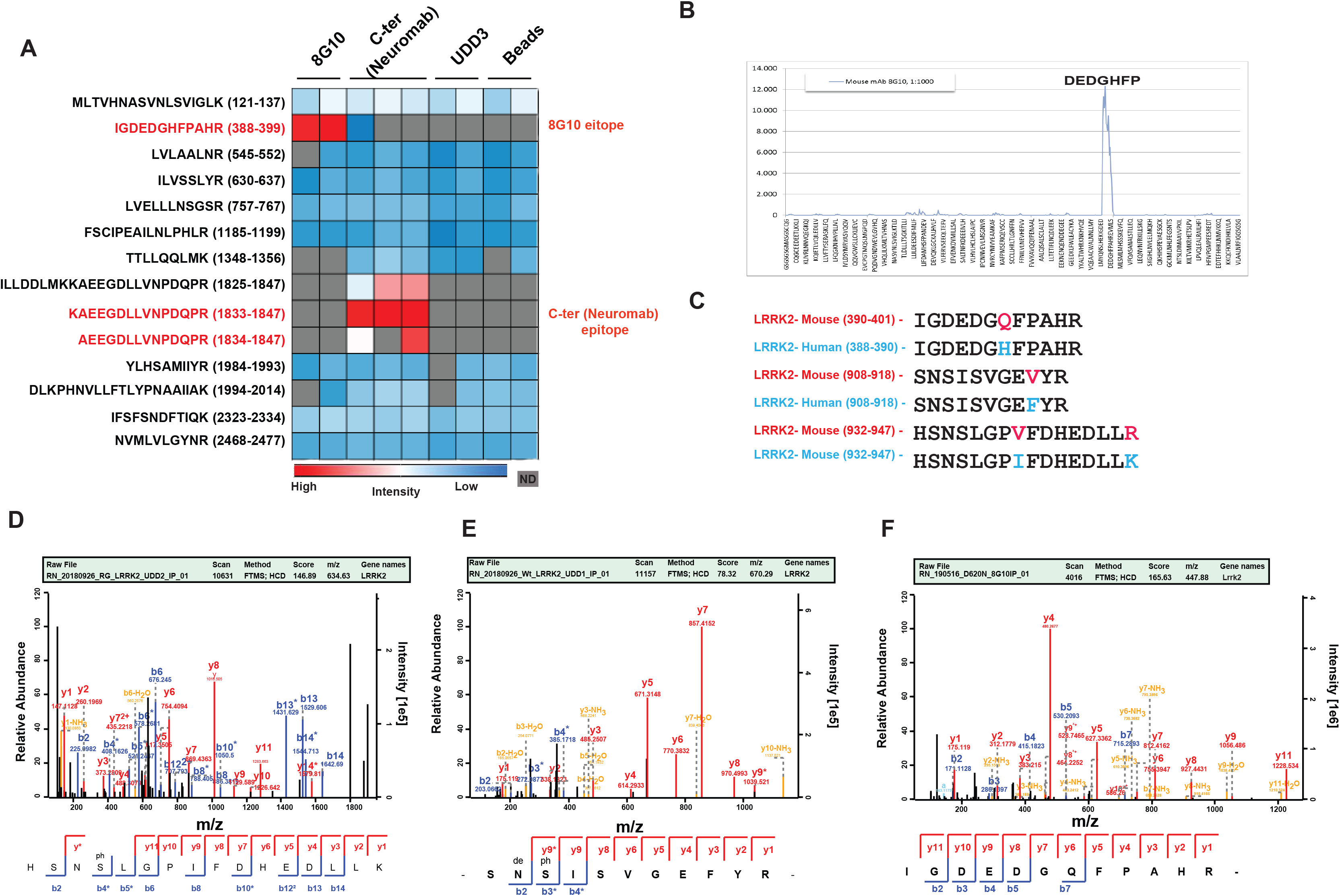
Epitope mapping of LRRK2 UDD1, UDD2, and LRRK2 C-terminal N241a/34 antibodies. **S3-A)** Wild type A549 cells were lysed, digested with trypsin and subjected to with the indicated antibodies. Eluted peptides were analysed by MS in data dependent mode and intensities of detected peptides represented on a heatmap blue (lowest) to red (highest). **S3-B)** an orthogonal validation of 8G10 antigen sequence by PEP preprint microarray-based technology. Custom prepared LRRK2 sequence peptide microarray was incubated with 1:1000 dilution of 8G10 antibody. Spot identification and quantification was carried out using LI-COR Odyssey imaging system. Highest intensity was observed for the LRRK2 sequence (DEDGHFP LRRK2 391-397). **S3-D)** MS/MS spectrum of an identified Phosphopeptide HSNSLGPIFDHEDLLK (LRRK2 pS935) that is selectively enriched by UDD2 antibody. **S3-E)** MS/MS spectrum of a Phosphopeptide SNSISVGEFYR (LRRK2 pS910) that is selectively enriched by UDD1 antibody. **S3-F)** MEFs, MS/MS spectrum of a mouse specific LRRK2 peptide IGDEDGQFPAHR (LRRK2 388-399) that was selectively enriched by 8G10 antibody in VPS35 [D620N] MEFs.

**Supplementary Figure 4:**
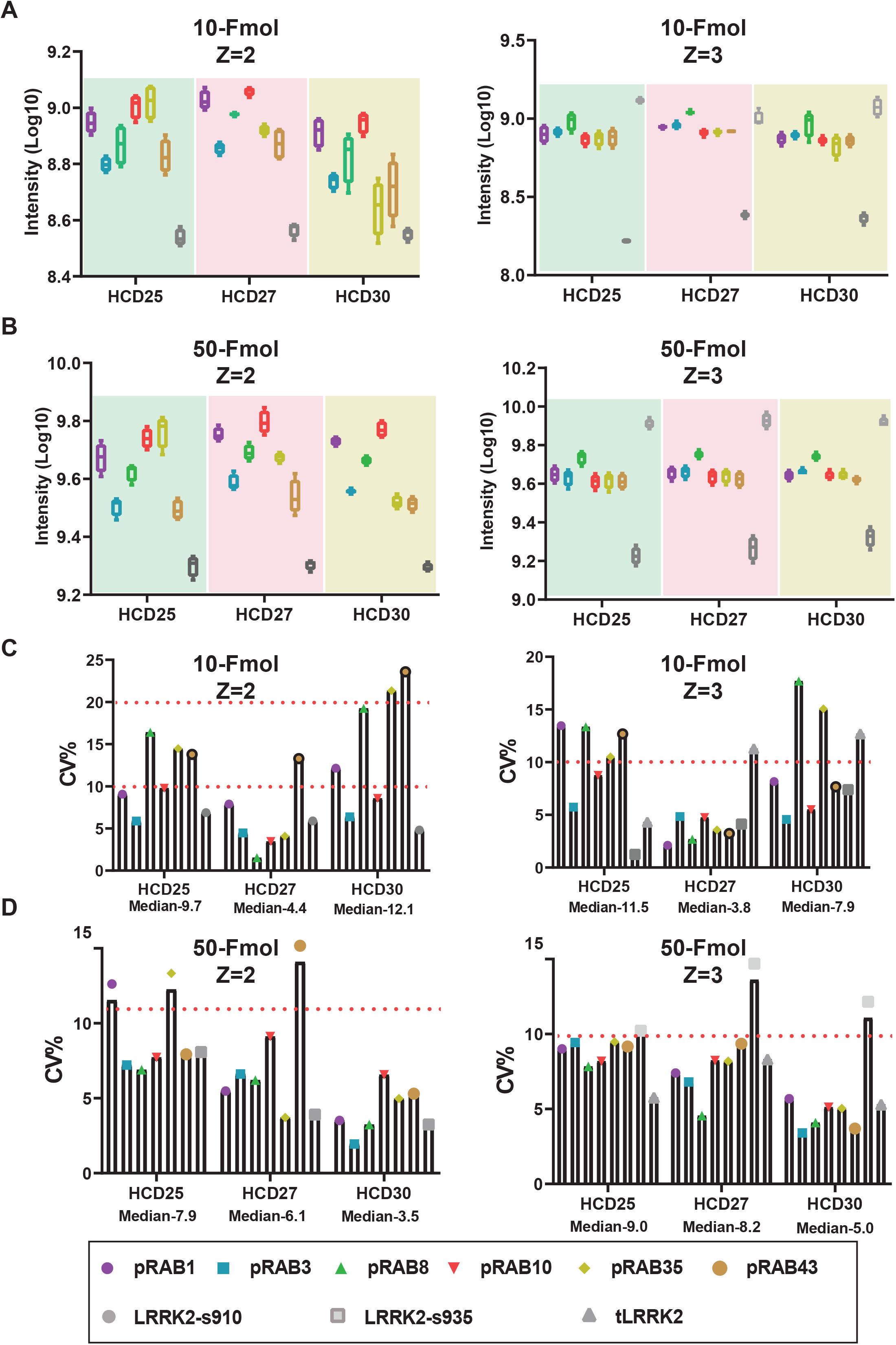
Determining the optimal collision energies for pRabs and pLRRK2 and total LRRK2 peptides: **S4: A-B)** Equimolar mixture of 10 fmol (S4-A) or 50 fmol (S4-B) containing six pRabs and LRRK2-pS910/pS935 and total LRRK2 were spiked into 50ng of HeLa peptide digest. HCD collision energies (CE) 25, 27 and 30 were tested to determine the best CE fragmentation for indicated peptides. Box plots depicting (S4: A-B) log10 intensities on y-axis and tested HCD CE value of X axis. **S4-C)** Bar graphs depicting the distribution of coefficient variation (CV) shown in percentage for 10 fmol amount experiments of all tested pRabs and pLRRK2 peptides on y-axis. X-axis indicating the median CV% value for HCD25, 27 and 30 respectively. **S4-D)** Bar graphs depicting the distribution of coefficient variation (CV) shown in percentage for 50 fmol amount experiments of all tested pRabs and pLRRK2 peptides on y-axis. X-axis indicating the median CV% value for HCD25, 27 and 30 respectively. n=3, error bars representing mean ±SEM

**Supplementary Figure 5:**
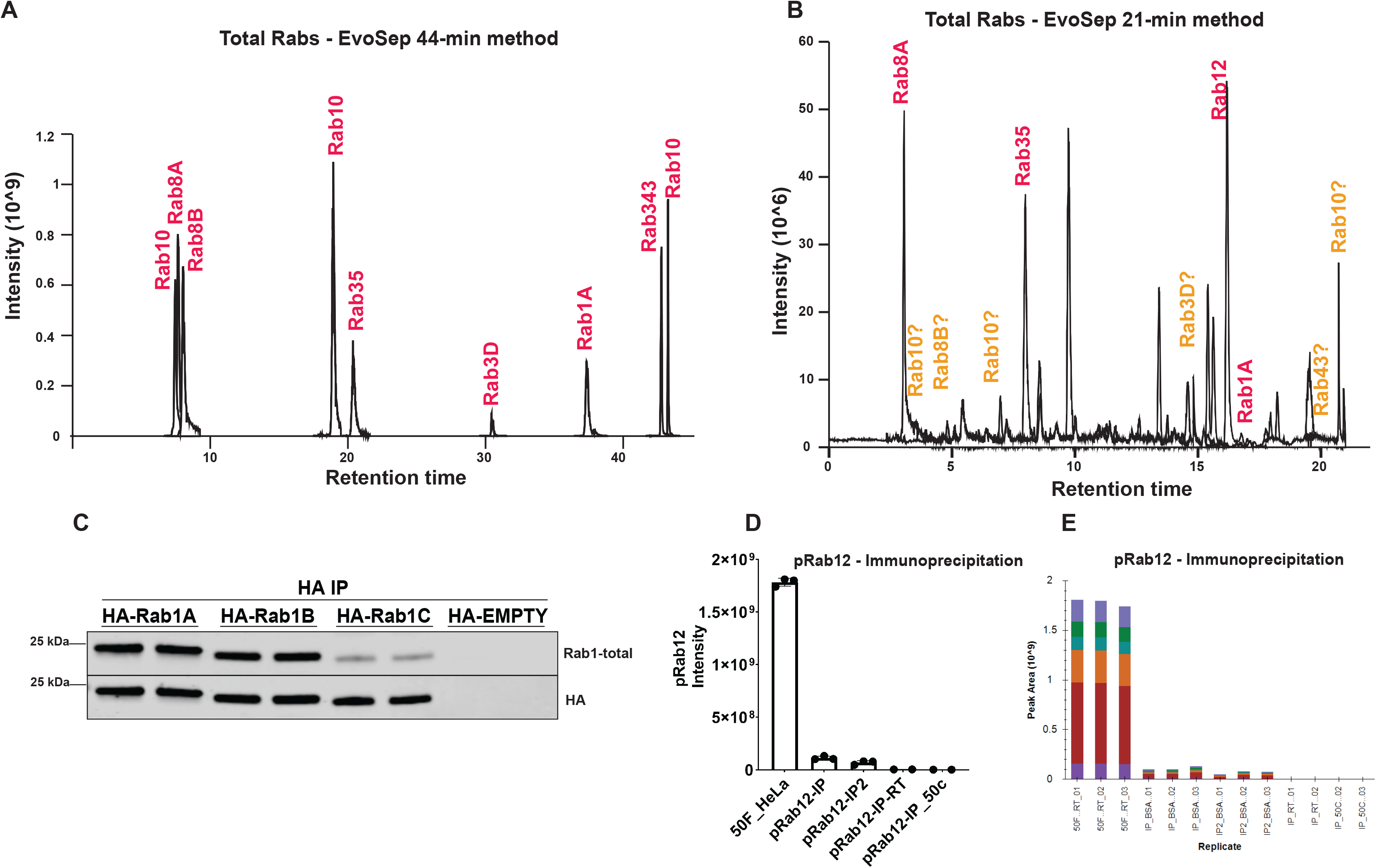
Development of total Rab proteins targeted PRM assay. S5-A) XIC depicting the 50 Fmol of an equimolar mixture of total Rab peptides (Rav1A, Rab1B, Rab3D, Rab8A, Rab8B, Rab10, Rab12, Rab35 and Rab43) was spiked into 50ng of HeLa and separated on EvoSep LC using 44 min. S5-B) XIC depicting 10 fmol of equimolar mixture of total Rab peptides separated on EvoSep LC using 21 min run. Retention times corresponding to each Rab was indicated in red coloured text (S5: A-B) and poorly separated Rab peptides in a 21 min run are indicated with orange coloured text. S5-C) Lysates of HEK293 cells overexpressing HA-Rab1A, HA-Rab1B, HA-Rab1C or HA-empty were used for HA immunoprecipitation (300 μg per IP). Immunoprecipitates were then immunoblotted with the Rab1-total and ant-HA antibodies. Immunoblots were subjected to quantitative LI-COR immunoblot analysis with all indicated antibodies at 1 μg/mL. S5: D-E) 50 fmol of heavy pRab12 spiked in 50ng of HeLa mixture was assessed to verify the immunoprecipitation efficiency using pRab12 antibody. 100Fmol of pRab12 heavy peptide was spiked into 200 μg of BSA tryptic digest and subjected to immunoprecipitation and analysed using PRM method. IP-2 indicating the flow-through was subjected to second round of Immunoprecipitation by adding fresh 1μg of pRab12 antibody. Immunoprecipitation was also assessed for samples Immunoprecipitated at room temperature and 50°C. S5-E) Peak areas depicting for the samples analysed in S5-C. n=3, error bars representing mean ±SEM

**Supplementary Figure 6:**
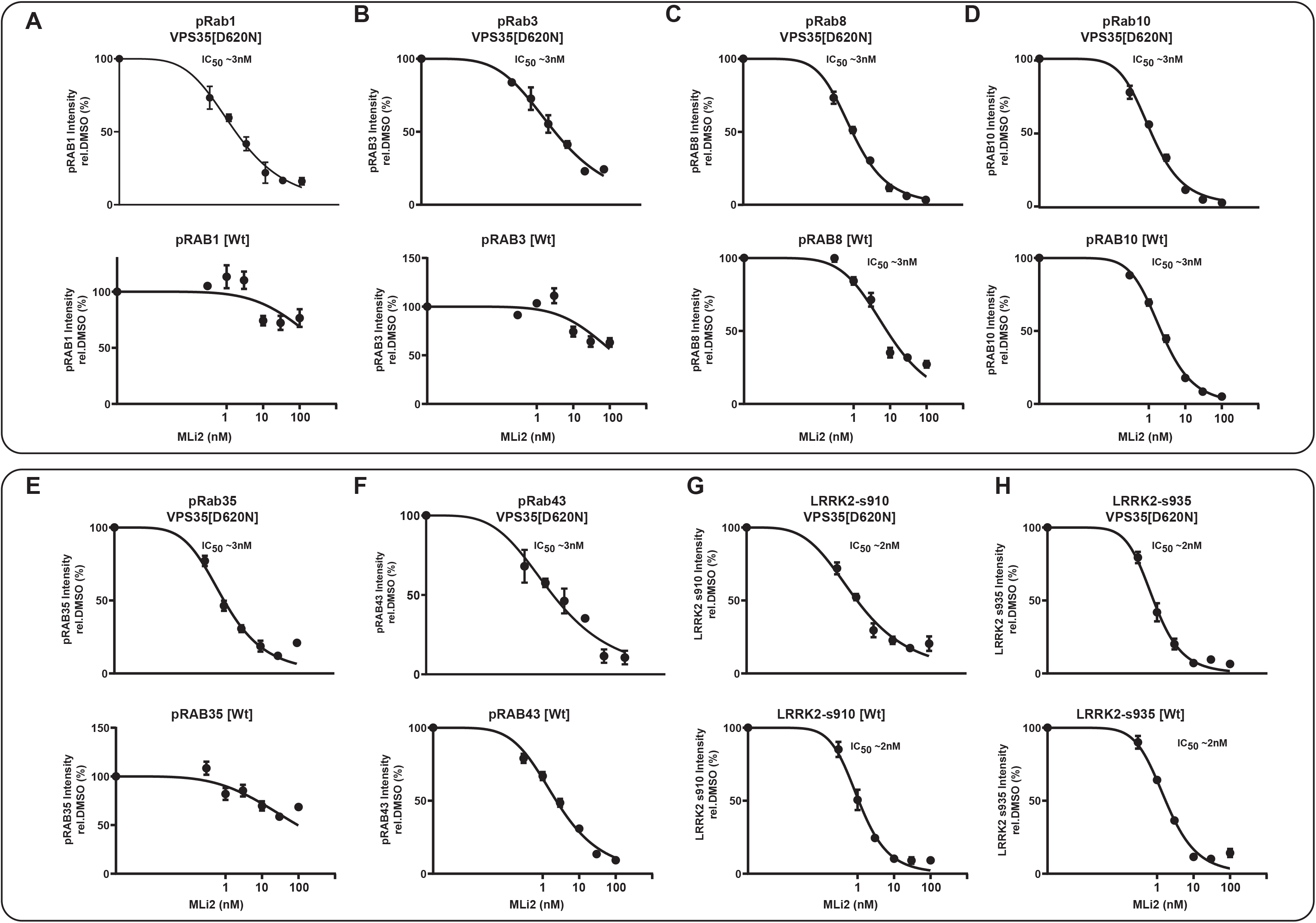
**(A-H).** MLi-2 inhibition IC50 values for both wild type and VPS35[D620N] were determined from the data presented in Figure 4, by dividing L/H ratio of each indicated MLi-2 concentration over the average value of L/H ratio of DMSO as 100%. The IC50 for each pRab was calculated using GraphPad prism software and indicated.

**Supplementary Figure 7:**
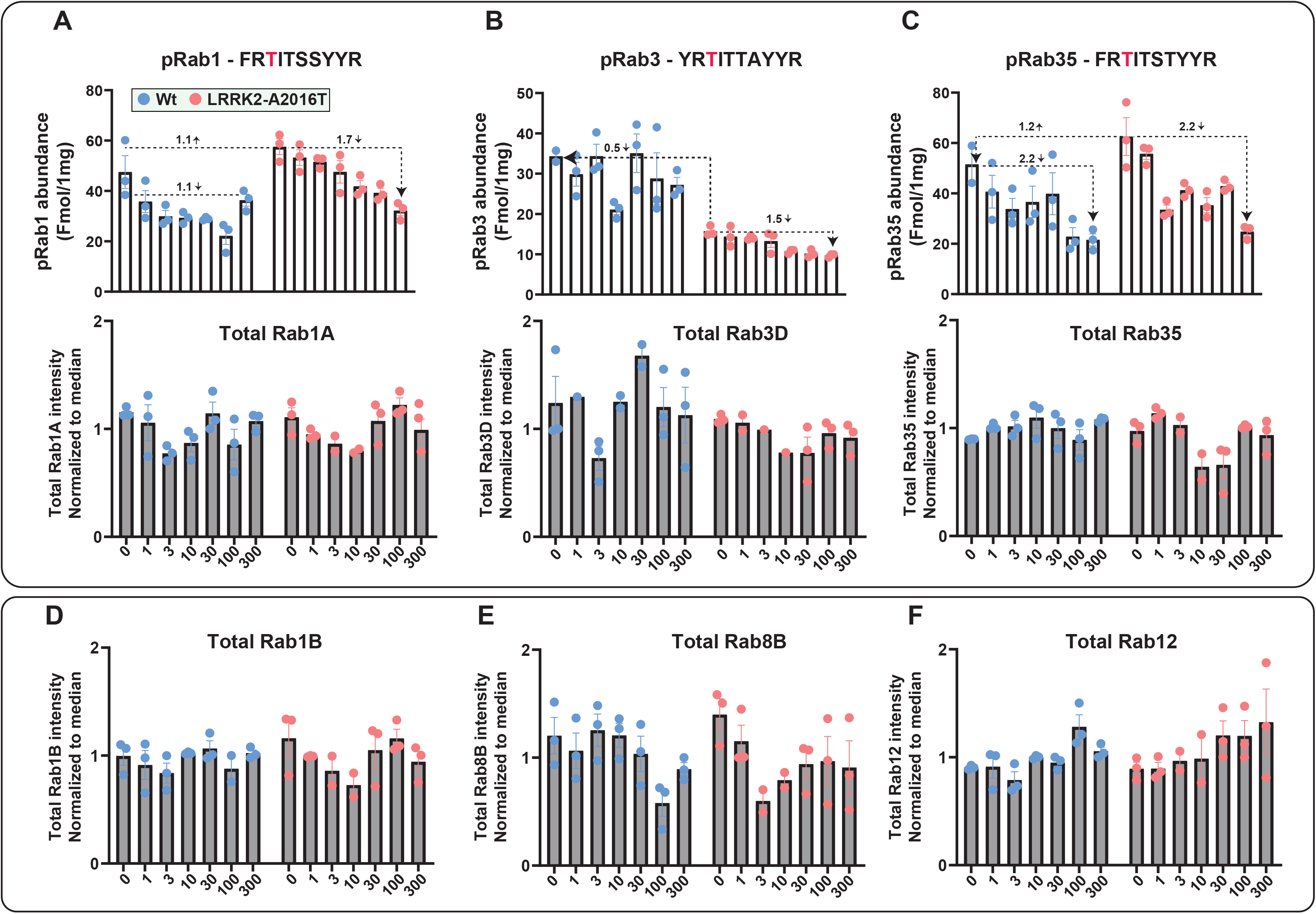
Data for pRab1, pRab3, and pRab35 derived from Figure 6.

**Supplementary Figure 8:**
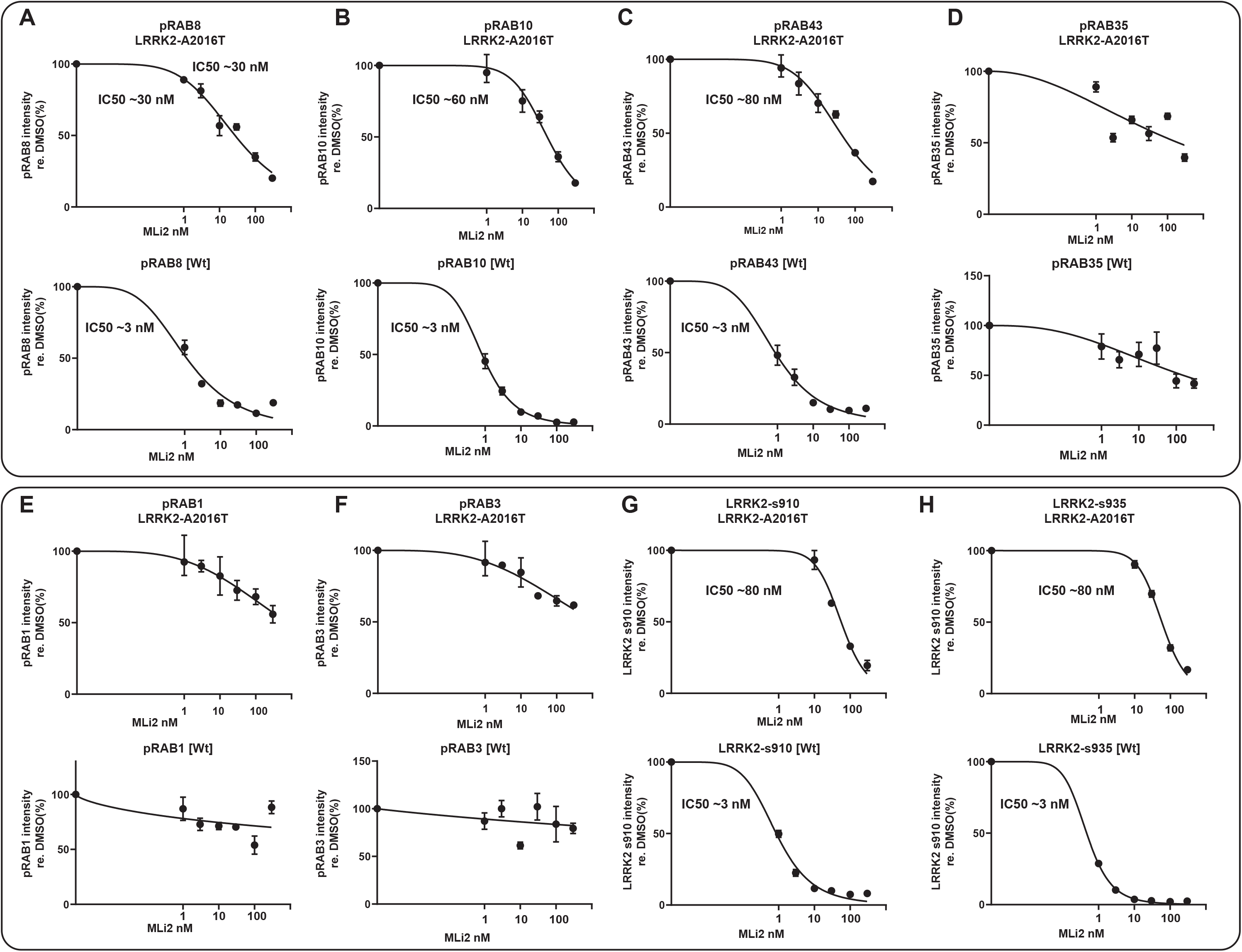
MLi-2 inhibition IC50 values for both wild type and LRRK2[A2016T] were determined from the data presented in Figure 5, by dividing L/H ratio of each indicated MLi-2 concentration over the average value of L/H ratio of DMSO as 100%. The IC50 for each pRab was calculated using GraphPad prism software and indicated.

**Supplementary Figure 9:**
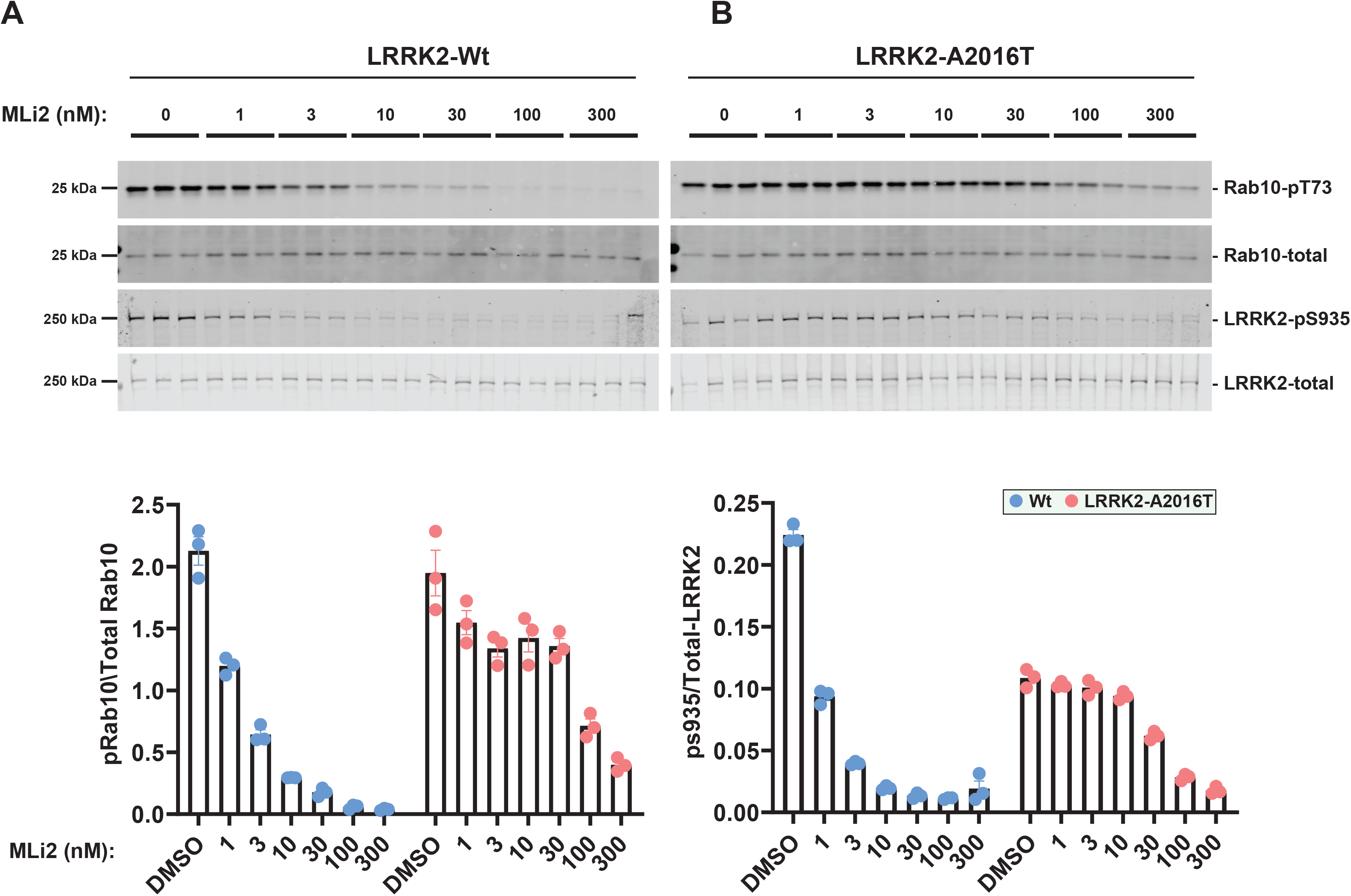
The indicated WT and LRRK2[A2016T] MEFs generated in Figure 5, were treated with or without the indicated concentrations of MLi-2 for 90 min. Cells were lysed, and 10 μg of extract was subjected to quantitative immunoblot analysis with the indicated antibodies (all at 1 μg/ml). Each lane represents cell extract obtained from a different dish of cells (3-4 replicates per condition). The membranes were developed using the Odyssey CLx Western Blot imaging system. Immunoblots were quantified using the Image Studio software. Data are presented relative to the phosphorylation ratio observed in WT cells treated with DMSO (no inhibitor), as mean ± SEM.

**Supplementary Figure 10:**
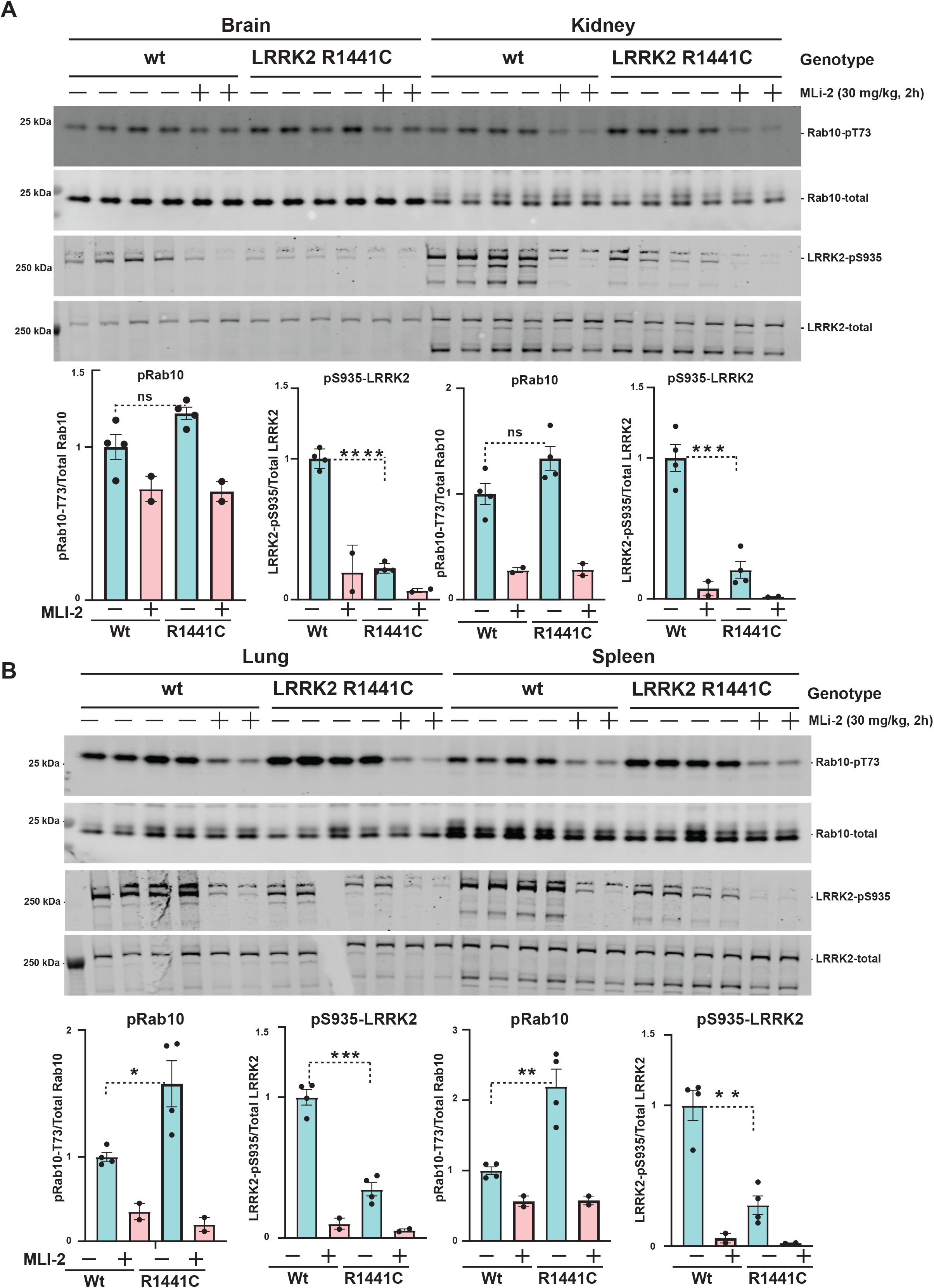
25 μg of extracts from the indicated tissues of WT and LRRK2 [R1441C] littermate mice harvested in Figure 8, were subjected to quantitative immunoblot analysis with the indicated antibodies (all at 1 μg/ml). Each lane represents cell extract obtained from a different animals (2-4 replicates per condition). The membranes were developed using the Odyssey CLx Western Blot imaging system. Immunoblots were quantified using the Image Studio software. Data are presented relative to the phosphorylation ratio observed in WT tissue (no ML-2). Statistical significance was determined by two tailed T-test and the significance of pRab10 (Brain: not significant P= 0.052), Kidney (ns P=0.067), Lungs (P=0.0207) and Spleen (P= 0.0015). For LRRK2 pS935 (Brain, (p=<0.00001), Kidney (P=0.0004), Lungs (P=0.000104) and Spleen (p= 0.001304). n=4 for vehicle and n=2 for MLi-2 treated mice. Error bars representing the mean ±SEM.

**Supplementary Figure 11:**
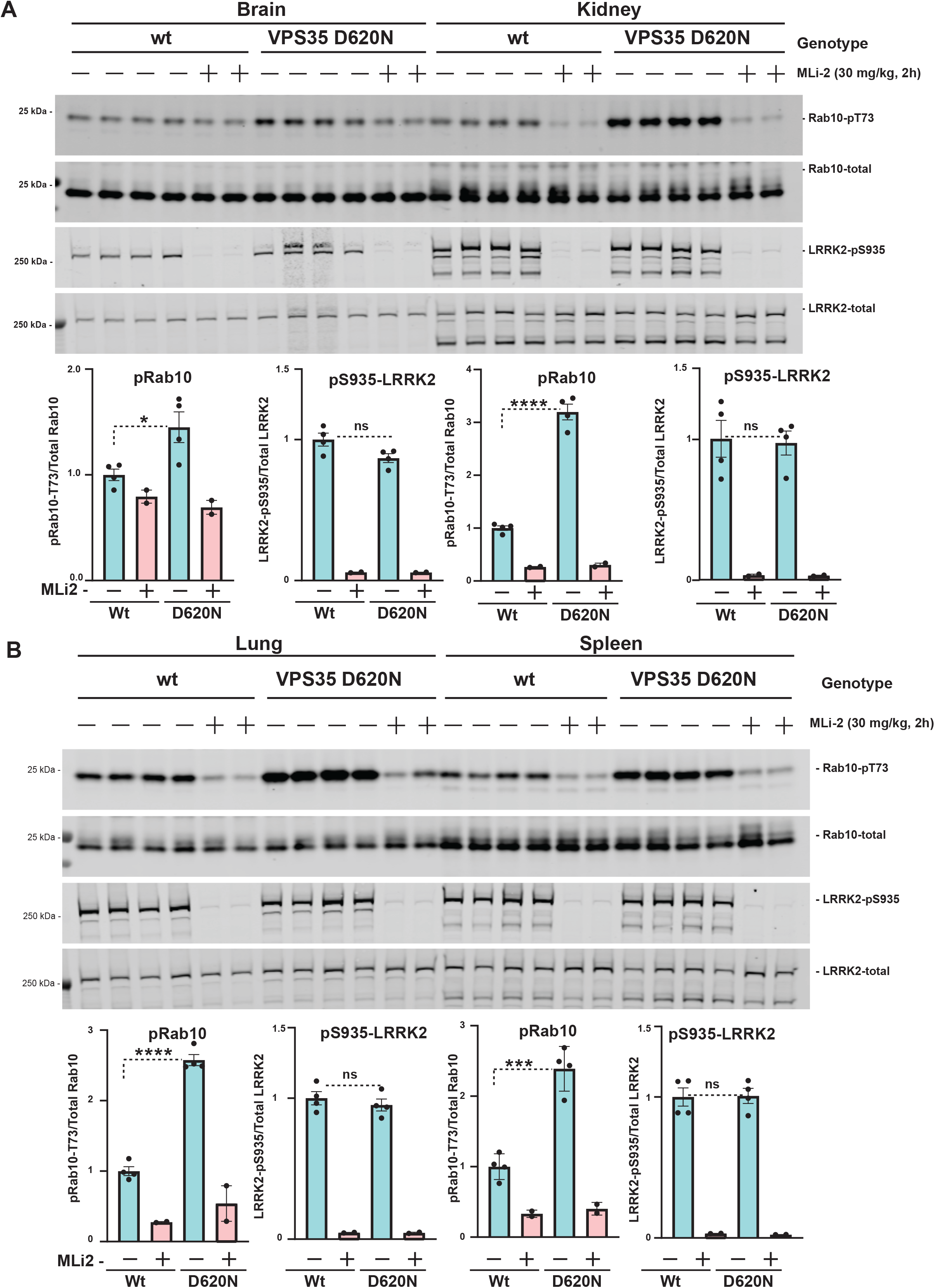
25 μg of extracts from the indicated tissues of WT and VPS35[D620N] littermate mice harvested in Figure 9, were subjected to quantitative immunoblot analysis with the indicated antibodies (all at 1 μg/ml). Each lane represents cell extract obtained from a different animals (2-4 replicates per condition). The membranes were developed using the Odyssey CLx Western Blot imaging system. Immunoblots were quantified using the Image Studio software. Data are presented relative to the phosphorylation ratio observed in WT tissue (no ML-2). Statistical significance was determined by two tailed T-test and the significance of pRab10 (Brain (P=0.0270), Kidney (P=<0.00001), Lungs (P=<0.00001) and Spleen (P=0.000271). For LRRK2 pS935 no statistical significance has been observed, (Brain, (p=0.47), Kidney (P=0.85), Lungs (P=0. 0.47) and Spleen (p= 0.93). n=4 for vehicle and n=2 for MLi-2 treated mice. Error bars representing the mean ±SEM.

**Supplementary Figure 12:**
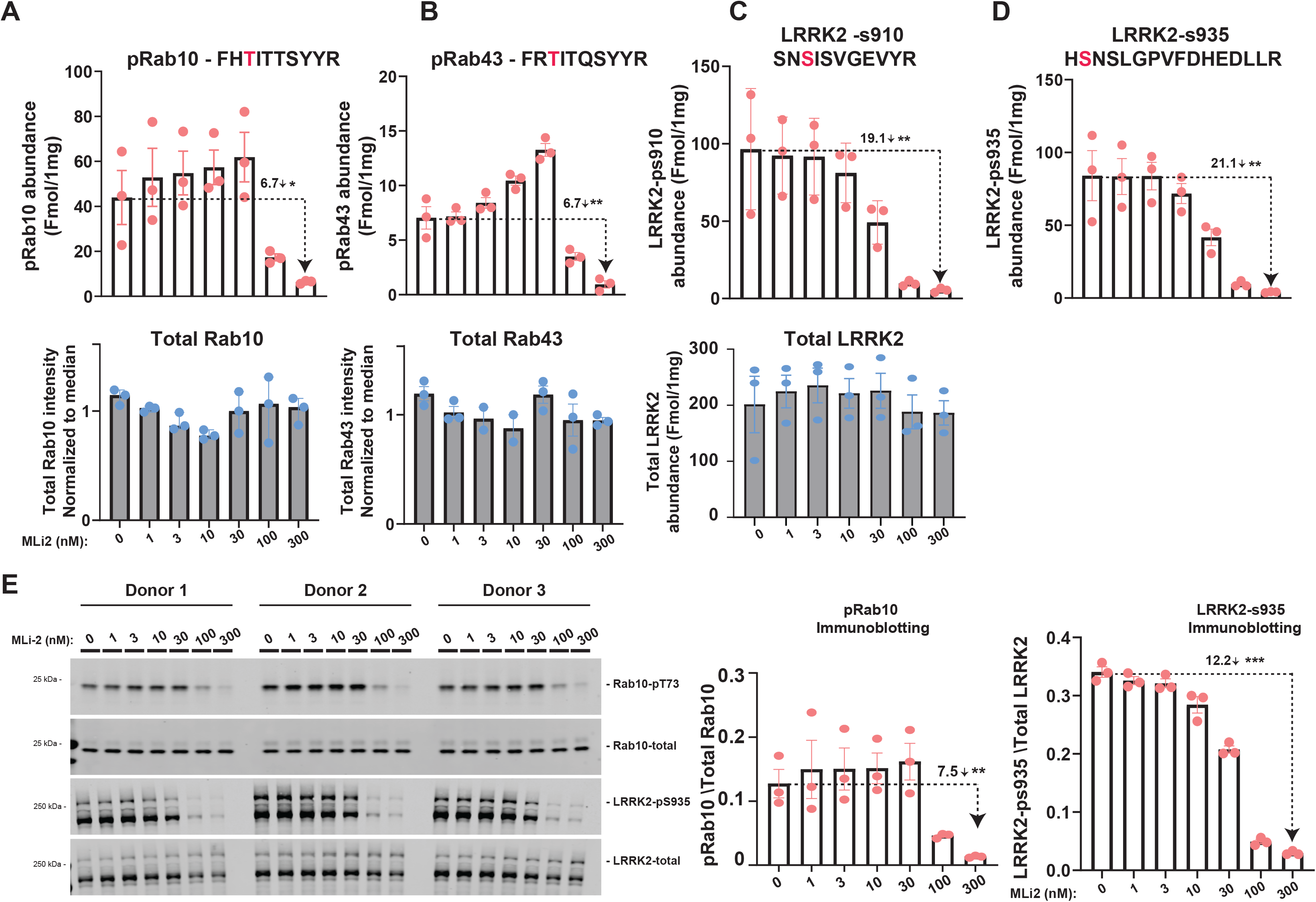
Targeted MS assay of LRRK2-Rab signalling pathway in human neutrophils. Primary neutrophils isolated from 3 healthy donors were treated with the indicated concentrations of MLi-2 for 60 min prior to harvest and subjected to either tryptic digestion (**A to D**) or immunoblot analysis (**E**). (**A to D**) 150 μg of digest was spiked with an equimolar ratio of 100 fmol heavy pRabs and LRRK2 pS910/pS935 and total LRRK2 peptides and subjected to sequential immunoprecipitation (IP) with a multiplexed antibody cocktail (Pan-pRab, UDD1, UDD2 and 8G10) and peptide levels quantified as fmol peptide/mg of tryptic digest (upper panel). In parallel, 0.5 μg of tryptic digest spiked with an equimolar ratio of 50 fmol of heavy total Rab peptides and data was acquired in PRM mode and peptide levels normalised to the median peptide intensity (lower panel). Dotted arrows indicated the differences between ± 300 nM MLi-2 samples and one tailed student t-Test was performed to depict the significance of change indicated with *. Three independent replicate experiments were performed for each of the indicated MLi-2 concentration and each dot indicates values for each experiment. Error bars representing mean ±SEM. (**E**) 10 μg of extract was subjected to quantitative immunoblot analysis with the indicated antibodies (all at 1 μg/ml). Each lane represents cell extract obtained from a different donors. Data are presented relative to the phosphorylation ratio observed in cells treated with DMSO (no inhibitor), as mean ± SEM. Statistical analysis was performed by one tailed student T-test indicating the significance between DMSO and 300nM MLi-2 conditions. A-D) Statistical significance represented in for PRM analysis are pRab10 (P= 0.017), pRab43 (P= 0.0024), LRRK2-pS910 (P=0.0078), LRRK2-pS935 (0.0048). G-F) Significance representing for the immunoblot analysis of pRab10 (p= 0.0034), LRRK2-pS935 (0.000556). Error bars representing the mean ±SEM.

**Supplemental table 1**: Details of reagents, antibodies, cDNA clones, Knock-in mice, MEFs, Heavy synthetic peptides equipment and software used in the current study.

**Supplemental table 2**: Identification and quantification of Wild type MEFs proteome, Wt VPS35 and VPS35[D620N] MEFs IC-50 pRabs PRM experiments, Wt LRRK2 and LRRK2[A2016T] MEFs IC-50 pRabs PRM experiments, Wt LRRK2 and LRRK2[R1441C] MEFs pRabs PRM experiments, Wt LRRK2 and LRRK2[R1441C] mice tissues (Whole brain, kidney, lungs and spleen) pRabs PRM experiments, Wt VPS35 and VPS35[D620N] mice tissues (Whole brain, kidney, lungs and spleen) pRabs PRM experiments, Human Neutrophils VPS35[D620N] patients and Healthy donors pRabs PRM experiments, Human Neutrophils IC50 pRabs PRM experiments and 21 min PRM QE HF-X instrument parameters used in this study.

**Supplemental table 3**: Identification and quantification of Wt VPS35 and VPS35[D620N] MEFs IC-50 total Rabs PRM experiments, Wt LRRK2 and LRRK2[A2016T] MEFs IC-50 total Rabs PRM experiments, Wt LRRK2 and LRRK2[R1441C] MEFs total Rabs PRM experiments, Wt LRRK2 and LRRK2[R1441C] mice tissues (Whole brain, kidney, lungs and spleen) total Rabs PRM experiments, Wt VPS35 and VPS35[D620N] mice tissues (Whole brain, kidney, lungs and spleen) total Rabs PRM experiments, Human Neutrophils VPS35[D620N] patients and Healthy donors total Rabs PRM experiments, HCD collision energy optimization experiments, pRab8 and pRab10 limit of detection experiments and 45 min PRM QE HF-X instrument parameters used in this study.

